# Shipped and shifted: modeling collection-induced bias in microbiome multi-omics using a tractable fermentation system

**DOI:** 10.1101/2025.08.15.670500

**Authors:** Annina R. Meyer, Jan Patrick Tan, Mihnea Paul Mihaila, Michelle Neugebauer, Laura Nyström, Nicholas A. Bokulich

## Abstract

Large-scale, decentralized microbiome sampling surveys and citizen science initiatives often require periods of storage at ambient temperature, potentially altering sample composition during collection and transport. We developed a generalizable framework to quantify and model these biases using sourdough as a tractable fermentation system, with samples subjected to controlled storage conditions (4 °C, 17 °C, 30 °C, regularly sampled up to 28 days). Machine-learning models paired with multi-omics profiling — including microbiome, targeted and untargeted metabolome profiling, and cultivation — revealed temperature-dependent shifts in bacterial community structure and metabolic profiles, while fungal communities remained stable. Storage induced ecological restructuring, marked by reduced network modularity and increased centrality of dominant taxa at higher temperatures. Notably, storage duration and temperature were strongly encoded in the multi-omics data, with temperature exerting a more pronounced influence than time. 24 of the top 25 predictors of storage condition were metabolites, underscoring functional layers as both sensitive to and informative of environmental exposure. These findings demonstrate that even short-term ambient storage (< 2 days) can substantially reshape microbiome, metabolome, and biochemical profiles, posing risks to data comparability in decentralized studies and emphasizing the need to recognize and address such biases. Critically, the high predictability of storage history offers a path toward bias detection and correction — particularly when standardized collection protocols are infeasible, as is common in decentralized sampling contexts. Our approach enables robust quantification and modeling of such storage effects across multi-omics datasets, unlocking more accurate interpretation of large-scale microbiome surveys.

## Introduction

Complex microbial ecosystems — microbiomes — colonize nearly every corner of our planet, playing key roles in food production as well as human, animal, plant, and planetary health (Banerjee et al., 2019; Berg et al., 2020; Gilbert et al., 2018). Accurate characterization of microbiomes and metabolome composition and function critically depends on standardized protocols for sample collection, storage, and handling. Even minor deviations in storage conditions can shift microbial community structure, altering metabolic outputs. These shifts can bias downstream analyses and compromise data comparability, as demonstrated in aquatic, soil, and gut microbiomes (Burman & Bengtsson-Palme, 2021); (Bokulich et al., 2019; McDonald et al., 2018; Scofield et al., 2015; Silva et al., 2022). While immediate freezing at -20°C or -80°C remains the gold standard to minimize microbial distortion in fecal microbiomes, as confirmed by both amplicon (Bassis et al., 2017; Tedjo et al., 2015, 2015) and whole metagenome sequencing (Momo Cabrera et al., 2025), it is often infeasible in decentralized, large-scale sampling efforts and in field studies. To limit compositional drift during room-temperature exposure, alternative stabilization strategies have been developed for fecal samples (Song et al., 2016), however, they are matrix-specific and do not generalize to diverse biological or food microbiomes.

This challenge is especially pronounced in citizen science and global-scale microbiome surveys, where samples are often shipped under heterogeneous ambient conditions without cold-chain logistics. In citizen science projects like the American Gut Project (McDonald et al., 2018), Global Sourdough Project (Landis et al., 2021), and HealthFerm Project (Meyer et al., 2025), samples frequently experienced uncontrolled temperature exposure before processing and analysis in the laboratory. Although storage effects on individual omics layers — such as microbial community composition — have been investigated, and correction strategies have been proposed for fecal samples (Amir et al., 2017; McDonald et al., 2018), the extent and nature of such distortions across integrated multi-omics data and in non-fecal matrices such as food fermentations remain underexplored.

In fermented foods, residual substrates sustain metabolism, enabling not only the outgrowth of individual taxa but also community-wide restructuring in response to environmental stressors (Marco et al., 2017; Sawant et al., 2025). Refrigeration slows, but does not halt these dynamics, and for example psychrotrophic bacteria such as *Hafnia alvei* and *Pseudomonas* spp. remain metabolically highly active even at 4 °C (Wei et al., 2019). Warmer storage accelerates metabolism and acidification, which initially contributes to food safety and spoilage prevention (Louw et al., 2023; Valentino et al., 2024). However, this can also accelerate community restructuring — particularly through the loss of lactic acid bacteria (LAB) viability (Cabello-Olmo et al., 2020; E. Kim et al., 2021; J. Y. Kim et al., 2022; Wei et al., 2019), favoring more acid- and heat-tolerant taxa (Louw et al., 2023). These shifts undermine both microbiome stability and also functional attributes of the fermentation ecosystems.

In parallel to taxonomic shifts, prolonged storage can induce changes in the metabolite landscape. Residual sugars and organic acids present in fresh fermentations are transformed by ongoing microbial metabolism, and can further reshape the biochemical fingerprint of a ferment before visible changes in microbial composition occur (De Filippis et al., 2016). Such transformations can obscure biological signals, compromise reproducibility, and confound interpretation in multi-omics studies.

Sourdough provides a tractable, ecologically relevant model to investigate these dynamics. Its microbiome, dominated by LAB, acetic acid bacteria, and yeasts, is highly responsive to environmental perturbations (Landis et al., 2021; Martins et al., 2021; Minervini et al., 2014; Sanmartin C, 2013; Van Kerrebroeck et al., 2017). Moreover, its low microbiome complexity makes it ideal for mechanistic studies of community and metabolic resilience.

While fermentation conditions have been extensively studied (De Vuyst et al., 2017; Ercolini et al., 2013; Reese et al., 2020; Ripari et al., 2016), the impact of post-sampling storage on sourdough microbiome and metabolome integrity remains poorly defined. Addressing this gap is critical — not only to safeguard interpretability in decentralized multi-omics studies of fermentations, but also to understand how environmental stress can reshape fermentation ecosystems, with potential implications for food quality and safety (Lim et al., 2024; Tan et al., 2022).

Here, we used sourdough as a tractable model fermentation system to investigate how realistic post-sampling storage conditions affect microbiome and metabolome profiles (Fig. 1). To simulate common sample shipping scenarios, sourdough aliquots were incubated at 4 °C, 17 °C, or 30 °C for up to 28 days without substrate replenishment. A temporal multi-omics design — spanning 16S rRNA and ITS amplicon sequencing, cultivation, untargeted FIA-MS metabolomics, and HPLC profiling — was used to dissect the progression of microbial and biochemical changes across storage conditions. Using machine learning, we demonstrated that both storage time and temperature were strongly and predictably encoded in multi-omics profiles, revealing consistent microbial and metabolic signatures of post-sampling environmental exposure. These patterns underscore the need for caution when interpreting data from decentralized sampling efforts, where storage conditions may confound biological signals.

**Figure 1.**
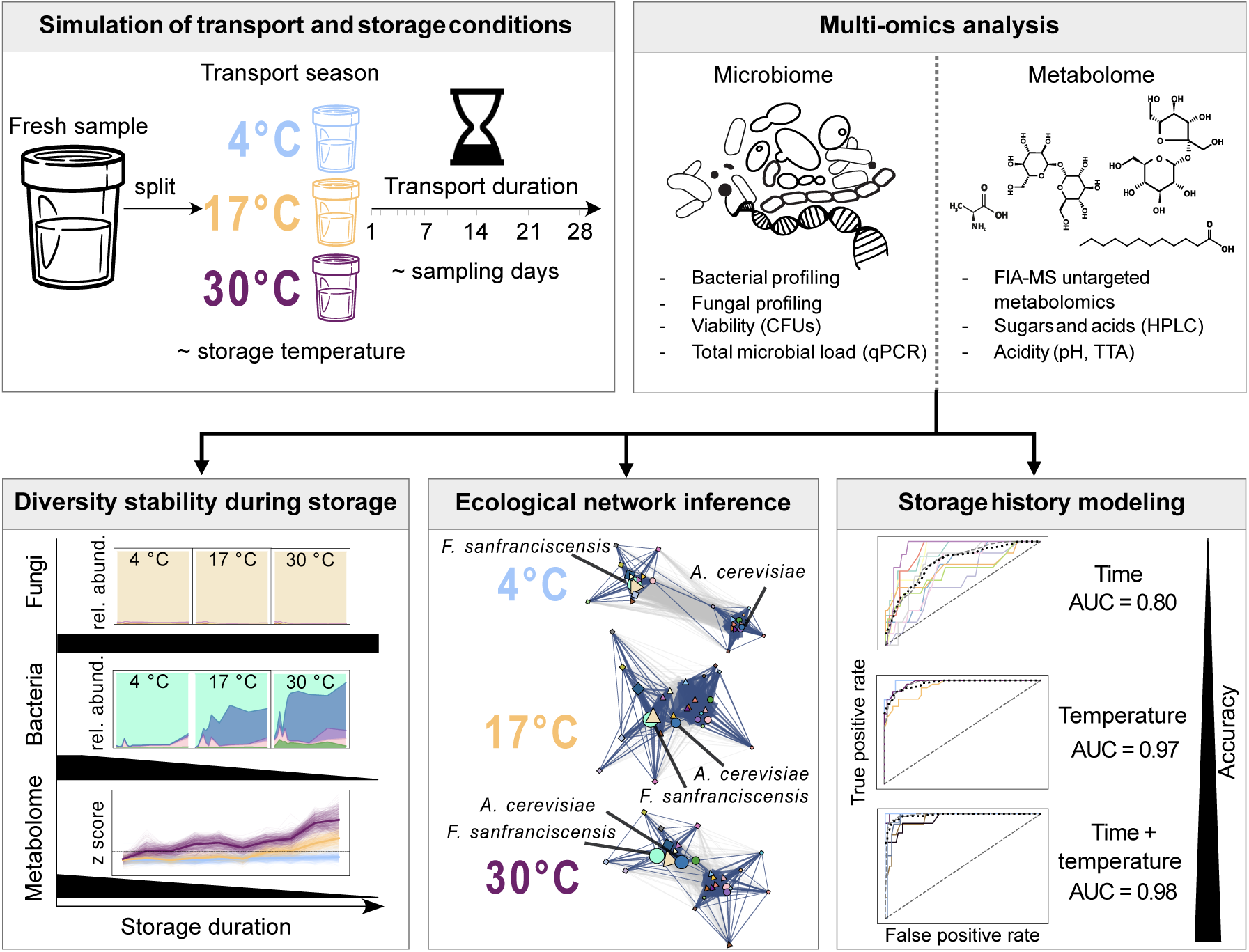
Overview of experimental design, multi-omics analysis, and main findings. A fresh sourdough starter was split and subjected to simulated post-collection storage scenarios across three temperatures (4 °C, 17 °C, and 30 °C), reflecting transport season. Samples were analyzed over 11 timepoints spanning 28 days, representative for variable transport durations (day 1-7, 10, 14, 21 and 28). A multi-omic approach was applied, combining microbiome (16S rRNA v4/ITS amplicon sequencing, qPCR, colony forming units (CFUs)) and metabolome (flow injection analysis mass spectrometry (FIA-MS), high-pressure liquid chromatography (HPLC), acidity) analysis to assess microbial and functional stability after variable collection conditions. Analyses revealed that bacterial diversity and metabolite profiles diverged significantly with prolonged warm storage, while fungal communities remained comparatively stable. Network inference highlighted ecological restructuring, notably the rise in centrality of the acetic acid bacterium *Acetobacter cerevisiae* with increasing temperature. Finally, supervised machine learning showed that integrated multi-omic features are sensitive predictors of storage time and temperature (AUC = 0.98), offering a basis for future bias detection and potential correction in decentralized microbiome studies.

## Results

### Temperature and time drive microbial viability loss and biochemical divergence in sourdough starters during simulated transport

Large-scale microbiome initiatives such as citizen science projects expand access to diverse ecosystems but often rely on ambient-temperature collection and transport, raising concerns about post-collection storage-induced bias. To quantify these effects under a controlled experimental setting, we simulated typical transport conditions using sourdough aliquots incubated at 4 °C, 17 °C, or 30 °C — representing seasonal ambient temperatures — over 28 days without substrate replenishment (Fig. 1).

At each time point, three biological replicates per temperature condition were analyzed. Microbiological analyses revealed rapid declines in viable lactic acid bacteria (LAB) and yeast at higher temperatures. By day 1, LAB colony forming units (CFUs) were already reduced at 30 °C relative to cooler conditions (Fig. 2a), with significant within-group declines from day 3 onward. Yeast viability declined from day 4–5 (Fig. 2b, Fig. S2c,d).

**Figure 2.**
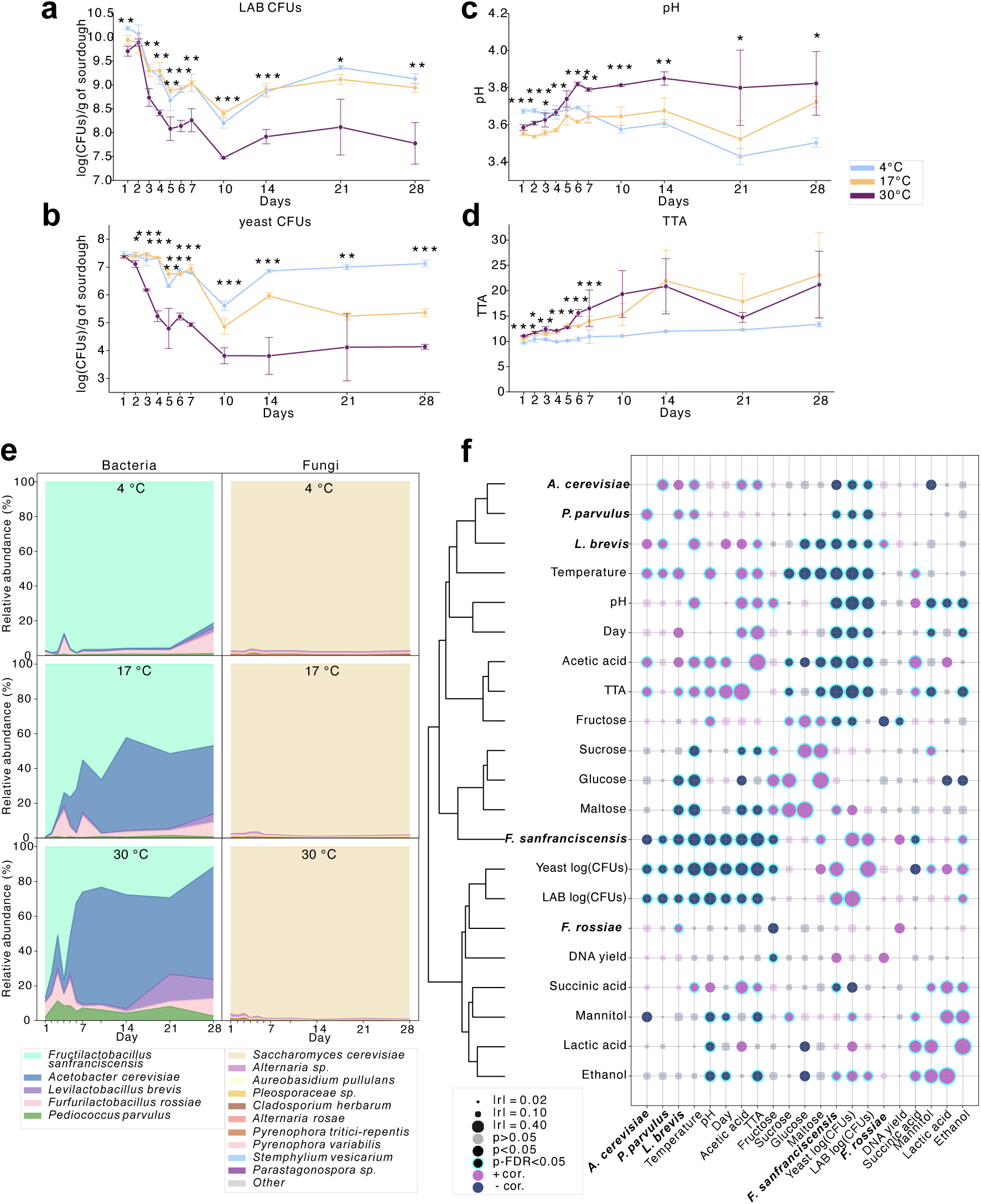
Simulated large-scale sourdough sampling reveals significant temperature- and time-dependent shifts in microbial and biochemical profiles. **a-d**, Temporal dynamics of lactic acid bacteria (LAB) CFUs, yeast CFUs, pH, and total titratable acidity (TTA). Statistical differences across temperatures at each time point were assessed by one-way ANOVA with FDR correction (*p < 0.05, **p < 0.01, ***p < 0.001). **e**, Relative abundance of bacterial (16S rRNA) and fungal (ITS) communities across storage conditions. Fungal taxa <0.05% were grouped as “Other”. **f**, Pearson correlation matrix of physicochemical and molecular parameters, including pH, TTA, CFUs, DNA yields, qPCR-based bacterial load, and HPLC-derived sugars and acids. Features were hierarchically clustered by pairwise correlations (r-values). All data represent biological triplicates.

Storage temperature also impacted acidity and metabolite dynamics. pH declined gradually at 4 °C but increased over time at 17 °C and 30 °C (Fig. 2c, Fig. S2a), while total titratable acidity (TTA) rose across all conditions, with a sharper increase at elevated temperatures — doubling by day 14 (Fig. 2d, Fig. S2b). These shifts reflect accelerated microbial metabolism under non-cooled conditions, further evidenced by high-pressure liquid chromatography (HPLC)-based metabolite profiling (Figs. S1–S3). Acetic acid showed accumulation at 17 °C and 30 °C from day six onwards (Fig. S1a), whereas lactic acid, succinic acid, and ethanol remained comparatively stable (Figs. S1b–d, S2e,f, S3a,b). Sugars such as glucose, maltose, and sucrose were rapidly depleted at warmer temperatures (Figs. S1e,g,h), followed by glucose reappearance from day 5–7 onward at 30 °C (Fig. S3c). Fructose accumulation from day 14 at 30 °C likely reflects enhanced enzymatic hydrolysis of oligosaccharides at elevated temperatures, potentially coupled with reduced microbial uptake or conversion (Fig. S1f). The concurrent stability of mannitol levels across all conditions (Fig. S1i, S3g) suggests a temperature-dependent suppression of fructose-to-mannitol conversion, which may be linked to the activity or abundance of mannitol-producing heterofermentative LAB.

### Higher temperature catalyzes microbial succession and promotes starvation tolerant taxa to bloom

Microbial community trajectories were profiled using 16S rRNA (V4) and ITS amplicon sequencing. The baseline bacterial community consisted of five species — *Fructilactobacillus sanfranciscensis*, *Acetobacter cerevisiae*, *Levilactobacillus brevis*, *Furfurilactobacillus rossiae*, and *Pediococcus parvulus* — while *Saccharomyces cerevisiae*, out of 27 detected fungal species, dominated the fungal community across all samples (>97.5% relative abundance; Fig. 2e).

At 4 °C, bacterial relative abundances remained stable throughout the one month period under starvation. However, warm conditions favored rapid expansion of *A. cerevisiae* (at 17 °C from day 3, and 30 °C from day 1), and also supported increased abundance of *L. brevis* and *P. parvulus*, while *F. sanfranciscensis* declined with increasing storage temperature and time (Fig. 2e).

To validate these trends and account for compositional bias, total bacterial load was quantified via 16S rRNA qPCR (Fig. S4a,d). Neither total abundance nor DNA yield varied significantly across time or temperature (Fig. S4b,c). Combining qPCR with relative abundance data enabled absolute quantification of species dynamics: *F. sanfranciscensis* remained dominant at 4 °C; at 17 °C, *A. cerevisiae* overtook by day 14; and at 30 °C, *A. cerevisiae* dominated by day 6 (Fig. S4e).

Correlational clustering of microbial, physicochemical, and metabolite data contextualized these shifts (Fig. 2f). *A. cerevisiae, P. parvulus*, and *L. brevis* co-clustered with temperature and pH, linking their activity to warming and acidification. This group was closely aligned with acetic acid, TTA, and fructose, indicating that elevated TTA is primarily driven by higher acetic acid accumulation relative to lactic acid. This is consistent with the higher pKₐ (≈ 4.75) of acetic acid compared to lactic acid (pKₐ ≈ 3.86). Conversely, F. sanfranciscensis clustered with readily fermentable sugars (glucose, maltose, sucrose), highlighting its more strict dependency on primary carbohydrate availability (Corsetti & Settanni, 2007). LAB and yeast CFUs, DNA yields, and residual sugars and acids formed a final cluster reflecting shared responses to broader ecological restructuring (Fig. 2f).

### Compositional restructuring hints at competitive dynamics under varying storage conditions

Alpha-diversity analyses reinforced the temperature- and time-driven bacterial succession observed in abundance profiles. Richness did not significantly change across any conditions for both bacteria and fungi, regardless of taxonomic resolution (ASV, OTU, k-mer; Fig. 3a,c; Fig. S5a,b,d). However, bacterial evenness increased significantly — by day 1 at 30 °C and from day 4 at 17 °C — indicating selective expansion of certain taxa under warmer conditions (Fig. 3b). Fungal evenness remained unchanged (Fig. 3d), consistent with *S. cerevisiae*’s near-total dominance across samples.

**Figure 3.**
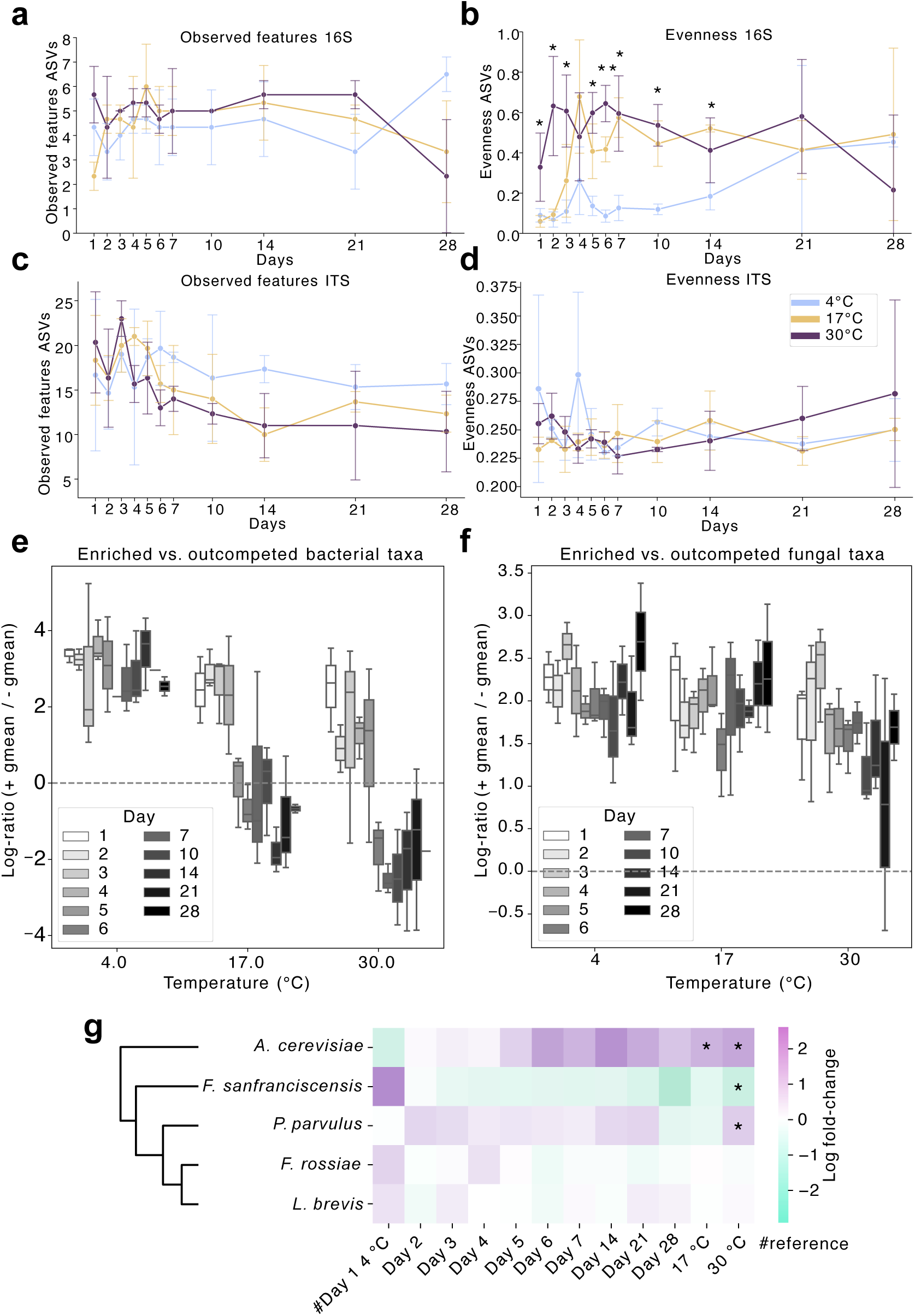
Storage temperature and duration drive bacterial, but not fungal, community restructuring in sourdough. **a–d**, Alpha-diversity of bacterial (16S; a, b) and fungal (ITS; c, d) communities over time and across storage temperatures. Richness (observed features) and evenness were calculated from ASV-level data; significance is assessed via one-way ANOVA with FDR correction (*p < 0.05, **p < 0.01, ***p < 0.001). **e–f**, Log-ratios of enriched versus outcompeted taxa based on reference group enrichments, with the reference for all comparisons being day 1 at 4 °C (ANCOM-BC2, Table S3 and S4), highlight pronounced bacterial compositional turnover at 17 °C and 30 °C, absent at 4 °C and across fungal communities. **g**, Differential abundance heatmap of bacterial taxa across conditions. Log-fold changes (relative to day 1 at 4 °C, again serving as reference for all comparisons across all temperatures and time points) were computed via ANCOM-BC2; taxa hierarchically clustered using average linkage (Euclidean distance). Asterisks indicate significant differential abundances (FDR-adjusted q < 0.05).

To capture compositional imbalance, we applied ANCOM-BC2 differential abundance modeling. This revealed sharp declines in the log-ratio of enriched to outcompeted bacterial taxa between days 4–5 at 17 °C and days 5–6 at 30 °C (Fig. 3e), reflecting a community-wide turnover. No such restructuring occurred at 4 °C or in fungal communities at any temperature (Fig. 3f; Fig. S6), highlighting bacterial communities’ higher sensitivity to storage induced metabolic stress in this model system. Strikingly, *A. cerevisiae* and *F. sanfranciscensis* exhibited strongly opposing trajectories across all timepoints and temperatures (Fig. 3g), supporting a model of contrasting ecological adaptations: *F. sanfranciscensis* thrives in cool, sugar-rich conditions, whereas *A. cerevisiae* is better adapted to warm, more nutrient-depleted and acidified environments.

### Temperature and time shape bacterial and fungal beta-diversity in opposite directions

To dissect microbial restructuring under storage, we analyzed beta-diversity in bacterial (Fig. 4a,b) and fungal (Fig. 4c,d) communities. Among bacteria, Bray–Curtis distances captured greater variance than Jaccard across all feature levels, with k-mer-based PCoA explaining the most (PCo1: 86.97%, PCo2: 6.54%), outperforming OTUs (79.44%/9.76%) and ASVs (78.0%/10.56%). This highlights the superior resolution of k-mers, even within a low-complexity system like sourdough.

**Figure 4.**
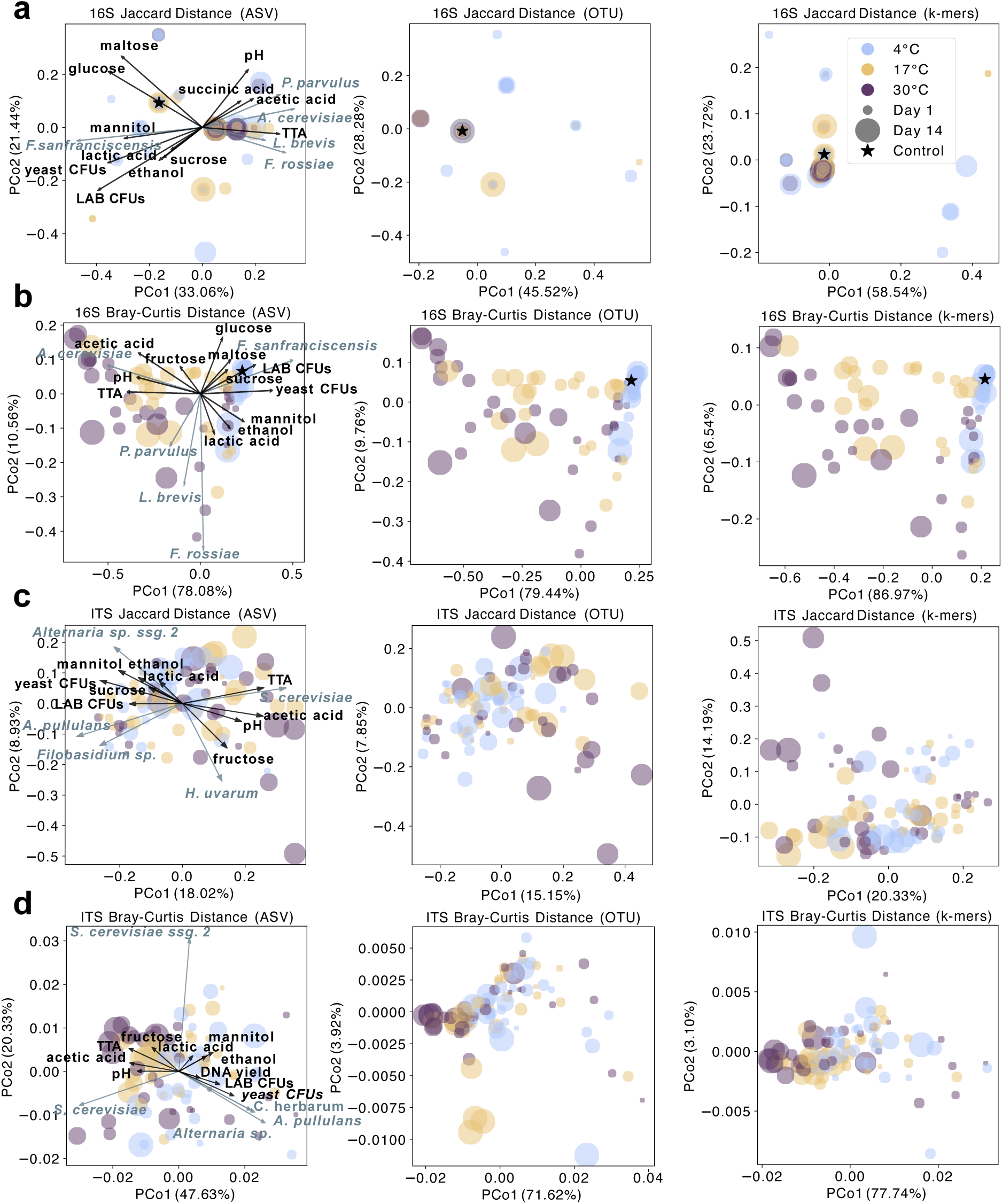
Storage modulates microbial community structure via compositional shifts, captured most sensitively by k-mer-based profiling. **a–d**, PCoA ordinations of bacterial (16S; **a–b**) and fungal (ITS; **c–d**) communities based on Jaccard (**a, c**) and Bray-Curtis (**b, d**) distance metrics across ASV, OTU, and k-mer resolutions. For ASV-based ordinations, the top taxa and metadata most correlated with the first two PCoA axes are overlaid as arrows, with direction and length reflecting Pearson’s r values. Arrows are scaled by the sum of correlations with PCo1 and PCo2. The ‘control’ refers to the mother sourdough at day 0 before splitting and incubating sourdoughs at either 4 °C, 17 °C or 30 °C; no ITS control is shown due to rarefaction-driven sample loss.

Biplot overlays revealed that 4 °C samples clustered with higher glucose, maltose, sucrose, mannitol, ethanol, lactic acid, CFU counts, and *F. sanfranciscensis*, while 17 °C and 30 °C samples correlated with *A. cerevisiae*, elevated acetic acid levels, TTA, pH, and fructose (Fig. 4b).

Fungal beta-diversity was less temperature-dependent, though Bray–Curtis again explained more variance. Two *S. cerevisiae* ASVs associated with high temperature and long storage correlated with TTA, pH, acetic acid, and fructose, while 4 °C samples were enriched in minor taxa, DNA yield, CFUs, mannitol, and ethanol (Fig. 4d).

Beta-dispersion analyses revealed rising bacterial heterogeneity with storage duration and temperature, as shown by increasing mean distances to centroid: 0.049 (4 °C), 0.240 (17 °C), 0.337 (30 °C), and over time from 0.064 (day 1) to 0.218 (day 4) and 0.356 (day 21). PERMDISP confirmed significant dispersion increases (temperature: F_ASV = 40.262, F_OTU = 46.166, F_kmer = 42.082; day: F_ASV = 3.508, F_OTU = 3.413, F_kmer = 4.160; all p = 0.001). In contrast, fungal dispersion remained stable (mean distance to centroid: 0.016–0.018; PERMDISP p > 0.2), indicating higher resilience to storage-induced stress.

PERMANOVA supported these patterns: for bacteria, variance was primarily explained by temperature (R² = 0.36, p = 0.001), followed by nested effects (temperature(day), R² = 0.35, p = 0.001), and day (R² = 0.14, p = 0.001) (Fig. S7a); for fungi, day had stronger effects (R² = 0.14, p = 0.001) than temperature (R² = 0.04, p = 0.009) (Fig. S7d). Nesting by temperature within day explained more variance than the inverse in both kingdoms (Fig. S7c,f).

Notably, nested PERMANOVA revealed divergent temperature-dependence: bacterial communities showed greatest temporal variance at 4 °C (R² = 5.168, p = 0.006), decreasing at 17 °C (R² = 4.284, p = 0.001) and 30 °C (R² = 3.164, p = 0.006); fungi displayed the opposite trend, with variance increasing from 4 °C (R² = 1.155, p = 0.320) to 30 °C (R² = 2.632, p = 0.007). All together, this suggests that while bacterial communities at higher temperatures became more compositionally variable, these changes were also more stochastic across replicates. Fungal communities, by contrast, exhibited stable dispersion but increasing PERMANOVA R² at higher temperatures, pointing to greater susceptibility to temporal restructuring as compared to bacterial communities, which was consistently shared across samples.

### Untargeted metabolomics reveals temperature- and time-dependent metabolite clusters with distinct functional enrichments

To expand beyond targeted chemical analysis, we applied untargeted FIA-MS metabolomics, detecting 1,097 ion features across all sourdough samples (Fig. S8). Elevated temperatures led to increased metabolic diversity: at 17 °C, the number of features doubled by day 7, and at 30 °C, this increase was already evident by day 2 compared to 4 °C and day 1 (Fig. S8a,c). In line with bacterial patterns, feature evenness remained stable at 4 °C but rose progressively at 17 °C and more rapidly at 30 °C (Fig. S8b,d).

Principal component analysis of FIA-MS profiles under different normalization schemes (TSS, log, z-score) revealed temperature-driven clustering analogous to 16S Bray–Curtis ordinations (Fig. 5a). TSS normalization provided the clearest separation (PC1 = 60.11%, PC2 = 21.54%), outperforming log (39.71%/5.26%) and z-score (51.89%/12.23%). Bi-plot overlays indicated strong correlations between 4 °C samples and fermentable sugars (sucrose, glucose, maltose), *F. sanfranciscensis*, CFUs, and ethanol. In contrast, warmer storage aligned with increased concentrations of acetic acid, pH, TTA, *A. cerevisiae*, and subdominant taxa (*P. parvulus, L. brevis, F. rossiae*) (Fig. 5a).

**Figure 5.**
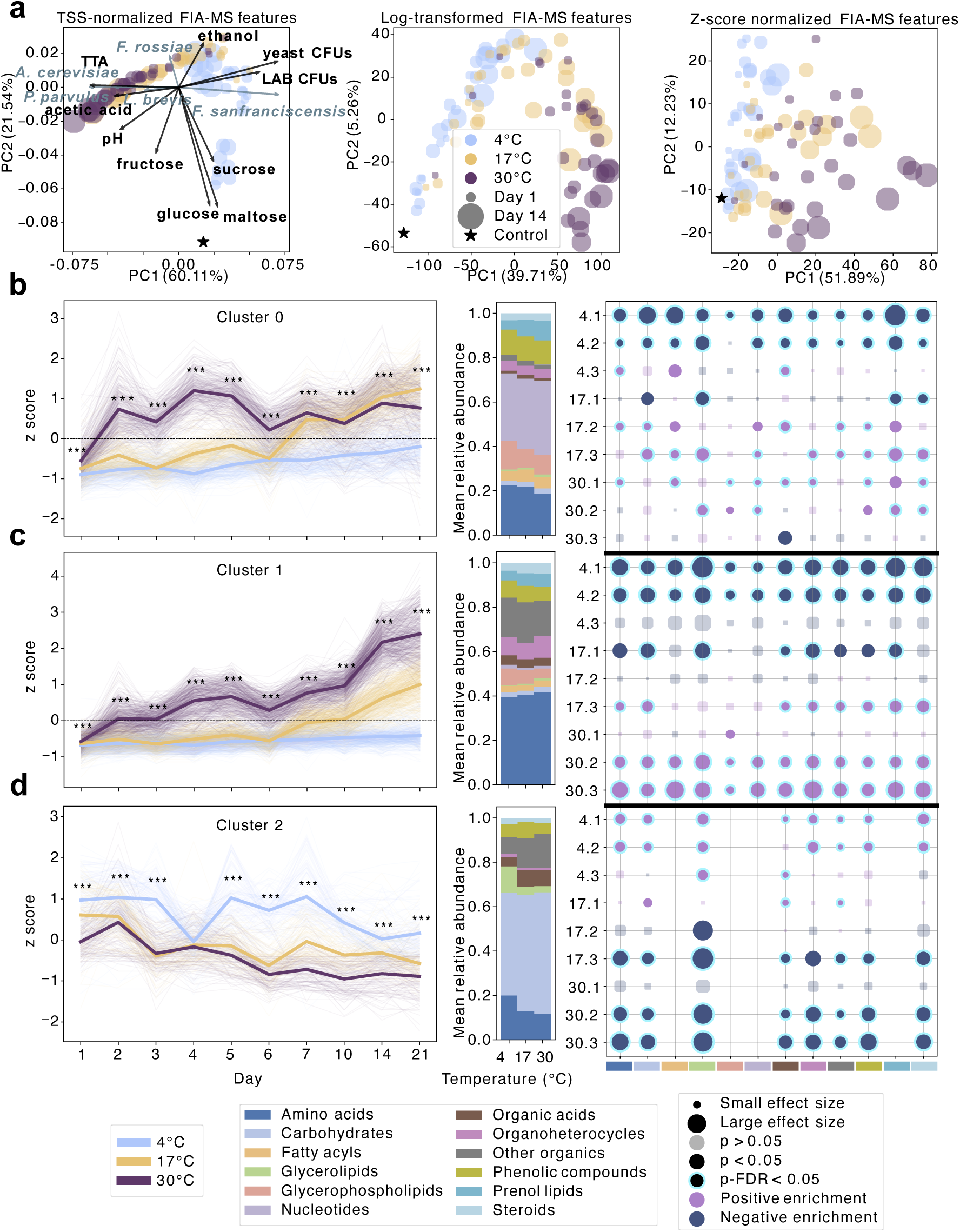
Storage conditions restructure the sourdough metabolome. **a**, Principal component analysis (PCA) of FIA-MS metabolite profiles under varying normalization and transformation strategies demonstrates how data preprocessing influences separation by temperature and explained variance. In the TSS-normalized ordination (left), key bacterial taxa and selected HPLC/metadata variables were projected as arrows. Arrow direction and length reflect Pearson correlations with PC1 and PC2, scaled to the sum of absolute correlations. **b–d**, Temperature- and time-responsive metabolites were clustered via k-means into three dynamic groups: Cluster 0 (b), Cluster 1 (c), and Cluster 2 (d). Z-scored metabolite trajectories are shown per temperature, with fine lines for individual features and bold lines for group means. Temperature effects at each time point were assessed via Kruskal–Wallis tests with FDR correction (*p < 0.05, **p < 0.01, ***p < 0.001). Bar plots show the average class-level metabolite abundances across temperature conditions. To identify associations between metabolite classes, time, and temperature, Mann– Whitney U tests were performed by comparing each feature in a given condition against all others. Time was binned into three phases: Phase 1 (days 1–4), Phase 2 (days 5–10), and Phase 3 (days 14–21). Each time phase was further stratified by temperature, with condition labels such as, e.g., 4.1 (Phase 1 at 4 °C) or 17.1 (Phase 1 at 17 °C). Enrichment was calculated as the log fold-change between condition-specific and global means. Statistically enriched metabolite classes (after BH-FDR correction) are annotated.

Procrustes analysis demonstrated strong concordance between metabolomic and bacterial community structures: FIA_TSS vs. 16S PCoA (M² = 0.437, p = 0.001), FIA_TSS PCA vs. 16S PCoA (M² = 0.498, p = 0.001), and FIA z-score vs. 16S (M² = 0.591, p = 0.001), with TSS offering the best alignment (Table S5). Concordance was highest at 17 °C (M² = 0.438), followed by 30 °C (M² = 0.593), and was non-significant at 4 °C (M² = 0.987, p = 0.718). Temporally, microbiome– metabolome synchrony peaked between days 6–14 (M² = 0.241–0.277, p < 0.01), diminishing during early and late storage phases (Table S6).

K-means clustering partitioned the FIA-MS features into three distinct temporal profiles (Fig. 5b– d), each reflecting coordinated biochemical responses (Kyoto Encyclopedia of Genes and Genomes (KEGG) + (Human Metabolome Database) HMDB annotation ± 0.002 Da, categorization into fermentation-relevant metabolite classes following the HMDB metabolite taxonomy).

- Cluster 0: Nucleotides, aromatic compounds, hormones, and toxin-like metabolites remained stable at 4 °C but increased at 17 °C and 30 °C (Fig. 5b), likely reflecting a combination of enhanced nucleotide turnover and passive release via cellular lysis under nutrient limitation and increased (auto-)acidification at elevated temperatures. Similar shifts in nucleotide pool shifts have been reported in lactic acid bacteria under nutrient limitation, acid, and oxidative stress which all stimulate increased nucleotide salvage, DNA/RNA repair, and the production of stress-signaling nucleotides such as c-di-AMP (Kilstrup et al., 2005; Papadimitriou et al., 2016).
- Cluster 1: Enriched in amino acids, fatty acyls, glycerolipids, organoheterocycles, prenol lipids, and steroids. These catabolic metabolites surged at 30 °C during days 5–10, appeared later at 17 °C, and remained depleted at 4 °C (Fig. 5c). These dynamics align with pronounced proteolysis, lipid remodeling, and stringent response mediated amino acid recycling under starvation stress and elevated temperatures (Papadimitriou et al., 2016). However, given the absence of significant total bacterial biomass accumulation (qPCR, Fig. S4e), non-microbial sources such as endogenous wheat proteases and lipoxygenases may also contribute, particularly in response to thermal activation.
- Cluster 2: Carbohydrate-related features (oligosaccharides, sugar alcohols) showed depletion at elevated temperatures and stability or slight enrichment in early phases at 4 °C (Fig. 5d), mirroring faster sugar consumption due to elevated enzymatic activity at warmer storage conditions with progressing time.

Together, these trajectories reflect a temperature-dependent modulation of metabolic activity, with carbohydrate catabolism occurring more rapidly from day 1 onwards at warmer temperatures, and more gradually at 4 °C. The subsequent accumulation of proteolytic and nucleotide-associated metabolites at 17 °C and 30 °C, and their delayed rise at 4 °C (e.g., Cluster 0), suggest that core metabolic processes proceed at different rates depending on storage temperature. Overall, the data support a model of progressive, rate-controlled metabolic layering, in which warmer conditions accelerate both the onset and magnitude of nutrient stress responses such as proteolysis and nucleotide turnover.

### Storage temperature restructures microbial-metabolite networks

To investigate how storage conditions reshape ecosystem connectivity, we constructed abundance-weighted co-correlation networks (Fig. 6a–c) and sparse conditional co-occurrence networks (Fig. 6d–f) for each storage temperature (4 °C, 17 °C, 30 °C). In the co-correlation networks, two major modules consistently emerged: the one containing dominant taxa and their metabolite partners, and the second one harboring subdominant species. At 4 °C, *F. sanfranciscensis, F. rossiae*, and *S. cerevisiae* dominated the core cluster (Fig. 6a). At 17 °C, network modularity diminished as clusters became more interconnected, with overall stronger co-correlations (Fig. 6b). By 30 °C, *A. cerevisiae* transitioned from the subdominant cluster (4 °C) to the inter-module bridge (17 °C) and ultimately joined the dominant hub (Fig. 6c). This trajectory was mirrored by increasing betweenness and eigenvector centrality for *A. cerevisiae*, and a concurrent decline in centrality for *F. sanfranciscensis* (Fig. S9).

**Figure 6.**
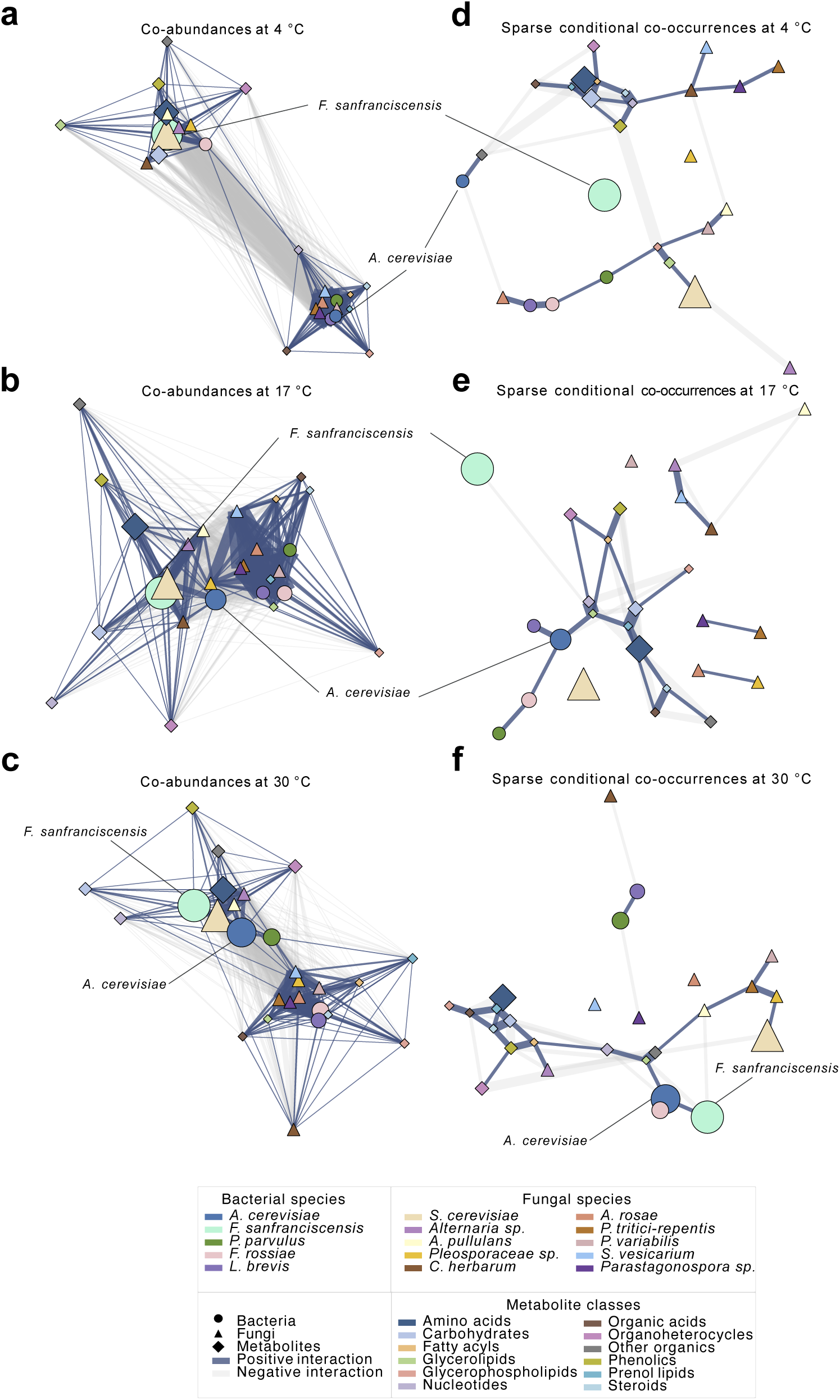
Ecological networks in sourdough microbiomes under starvation at different temperatures. **a–c**, Abundance weighted co-occurrence networks (CLR-transformed) reveal temperature-dependent microbial–metabolite association modules. At 4 °C, *F. sanfranciscensis* and yeast taxa dominate, while at 17 °C and especially 30 °C, *A. cerevisiae* becomes central, bridging core and subdominant taxa. **d–f**, Sparse inverse-covariance networks capture conditional dependencies: at low temperature, co-occurrences are compartmentalized; at higher temperatures, *A. cerevisiae* is a clear keystone, showing direct conditional links to both bacterial and metabolite nodes.

Sparse conditional networks uncovered temperature-specific rewiring of direct associations. At 4 °C, *F. sanfranciscensis* remained largely disconnected, with no significant positive microbiome or metabolome associations, while less dominant bacterial species exhibited stronger connectivity (Fig. 6d). Centrality analyses confirmed *F. sanfranciscensis* had the lowest centrality values at 4 °C (Fig. S10), whereas *A. cerevisiae* retained higher influence across all temperatures. At 30 °C, *F. sanfranciscensis* gained closeness centrality and established a positive edge with *A. cerevisiae* (Fig. 6f), suggesting increased ecological interdependence in the bacterial sourdough community under warmer fermentation conditions.

Together, these network analyses reveal that rising temperatures not only shift microbial abundances but also rewire co-occurrence topology - amplifying the integrative role of *A. cerevisiae* while diminishing the dominance of *F. sanfranciscensis*. This rewiring may reflect broader ecological adaptation across microbial and metabolic layers.

### Functional metabolite profiles outperform taxonomic composition in predicting storage history

To identify which features most robustly reflect storage-induced perturbations, we benchmarked predictive performance of individual and integrated data layers — including amplicon-based microbiome profiles, untargeted FIA-MS metabolomics, and physicochemical metadata (HPLC sugars/acids, CFUs, pH, TTA) — using cross-validated random forest models (Fig. 7a, S11).

**Figure 7.**
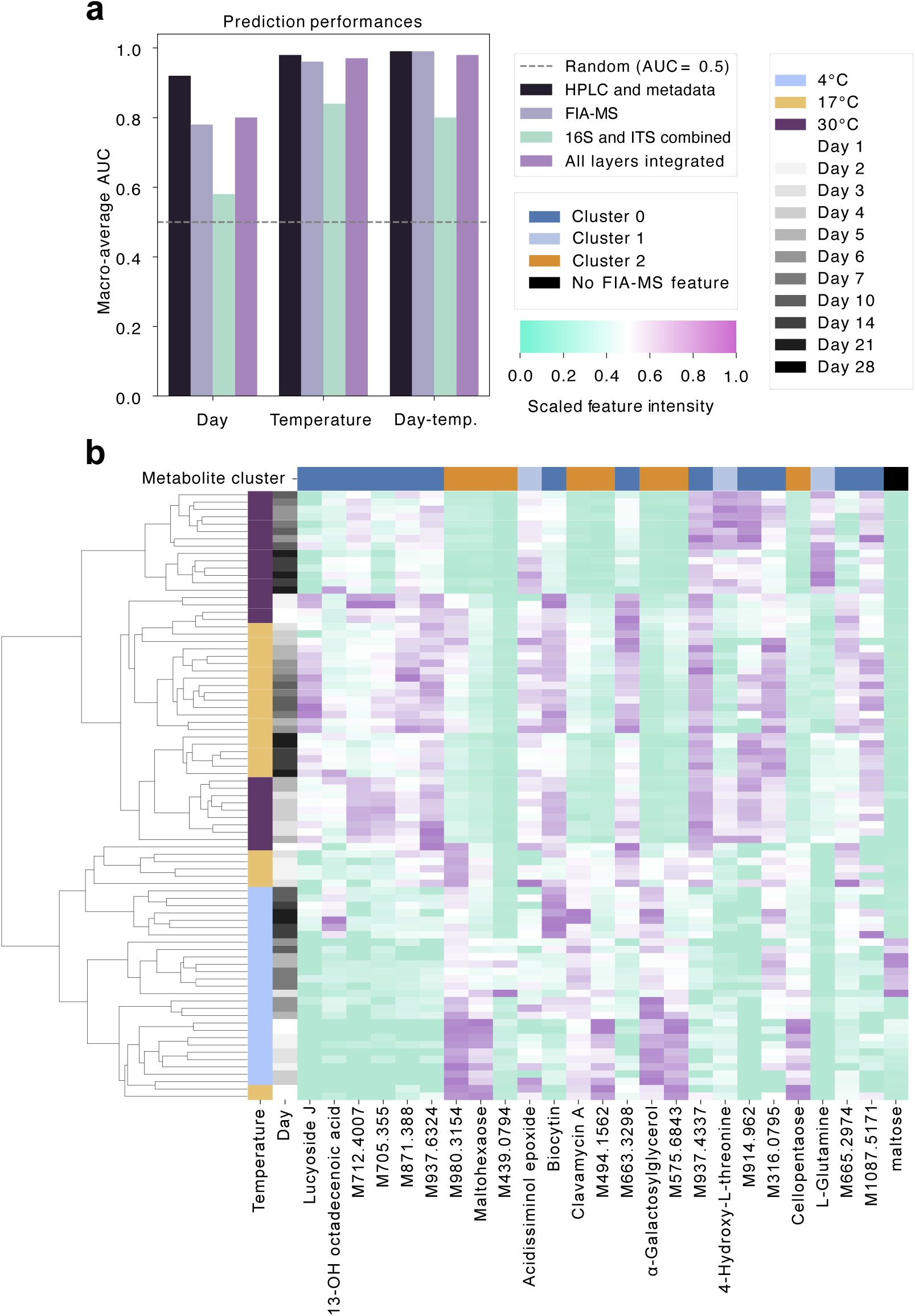
Storage-phase classification and predictive markers in sourdough microbiomes. **a**, Accuracy of the predictive performance for storage day, temperature, and combined day– temperature categories based on different data types: HPLC and metadata (including CFU counts, pH, and TTA), full FIA–MS metabolome profiles (annotated and unannotated), combined 16S and ITS microbiome data, and the fully integrated multi-omics dataset. Predictions were generated using cross-validated random forest (RF) models. **b**, Top 25 predictive features: strongest signals derive from metabolomics (Clusters 0–2), complemented by HPLC-measured sugars and acids and CFUs, the heatmap shows hierarchical clustering by storage phase (time x temperature) across metabolite clusters.

Storage duration was most accurately predicted from HPLC and metadata (macro-average AUC = 0.92), with LAB CFUs, fructose, and mannitol as top-ranked features (Fig. S12a). Predictive performance for storage duration based on the microbiome was poor (AUC = 0.58), and driven by low-abundance fungi (e.g., *P. variabilis*, *D. fristingensis*, *A. pullulans*) and *A. cerevisiae* (Fig. S13a), highlighting limited temporal specificity in community composition alone. Storage temperature was again best captured by HPLC features and metadata (AUC = 0.98), dominated by maltose, glucose, and TTA (Fig. S12b). The microbiome was more predictive for temperature (AUC = 0.84) than time, where most discriminatory features included *F. sanfranciscensis*, *P. parvulus*, and *P. variabilis* (Fig. S13b). This likely reflects their differential resilience to temperature-induced accelerated acidification, contributing to temperature-dependent niche structuring and community dynamics (Minervini et al., 2014; Papadimitriou et al., 2016).

Classification of combined temperature–time phases (Phase 1: days 1–4, Phase 2: days 5–10, Phase 3: days 14–28) achieved near-perfect accuracy with FIA-MS (AUC = 0.99), HPLC + metadata (AUC = 0.99), and full multi-omics integration (AUC = 0.98). Using microbiome data alone demonstrated high accuracy but inferior performance (AUC = 0.80) (Fig. S11). Strikingly, 24 of the top 25 predictive features originated from the metabolome, particularly FIA-MS Cluster 0, enriched in nucleotides, steroids, and stress-response metabolites with increasing abundance under warmer conditions and extended storage (Fig. 7b, S13f). Additional contributions came from Clusters 1 (amino acids) and 2 (carbohydrates), reflecting metabolic transitions from carbohydrate depletion to proteolysis and secondary metabolite accumulation. HPLC-measured maltose was the only non-FIA-MS predictor in the top 25.

These results highlight that metabolite profiles — reflecting the functional output of microbial activity — more robustly capture storage history than taxonomic composition alone. This decoupling of function from community structure underscores the central role of metabolic adaptation in driving fermentation dynamics, particularly under varying temperature conditions and extended storage durations.

## Discussion

Decentralized microbiome sampling — crucial to citizen science and global biodiversity surveys — faces a persistent challenge: preserving data integrity in the absence of controlled post-collection storage. Understanding how environmental exposures can reshape microbial composition and function is therefore critical for elucidating ecological dynamics and eliminating bias in large-scale microbiome studies. Our multi-omics analysis revealed that even short-term ambient storage (17–30 °C) restructures microbial ecosystems across multiple omics layers. In a controlled sourdough model, bacterial communities, metabolite profiles, and ecological network topology underwent pronounced shifts with storage temperature and duration — while fungal communities remained stable. These results highlight the higher sensitivity of bacterial and functional layers to environmental perturbation as compared to the fungal community, as observed in our relatively low-complexity, closed sourdough ecosystems, and can be modeled with high accuracy. More broadly, they reinforce that post-collection storage is not a neutral holding phase, but a dynamic window of ecological change with profound implications for microbiome data integrity and implications for reproducibility, comparability and multi-omic data interpretation.

Importantly, shifts in metabolome profiles preceded microbial community restructuring, and temperature — not time — was the primary lever. In the sourdough model at 4 °C, the core sourdough bacterium *Fructilactobacillus sanfranciscensis* remained dominant throughout, reflecting its cold tolerance and efficient sugar utilization in mildly acidic environments (Corsetti & Settanni, 2007), and metabolite profiles remained relatively stable over time. In contrast, higher temperatures accelerated substrate depletion — particularly maltose and sucrose — triggering a succession from LAB to more acid- and storage stress-tolerant taxa such as *Acetobacter cerevisiae*, consistent with their known higher acid tolerance and metabolic flexibility (Baig et al., 2021; Gomes et al., 2018; Han et al., 2024; Wang et al., 2015). This trajectory exemplifies an environmental filtering process (Marsland et al., 2019), wherein acid accumulation and resource scarcity seem to shift the community from sourdough specialization toward resilience and higher generalism. In line with this, *F. sanfranciscensis*, with its smaller genome and narrower metabolic repertoire (Vogel et al., 2011), declined with increasing storage temperature and time (Fig. 2e).

These observed compositional transitions were likely driven by a combination of ecological stressors and metabolic interdependencies. As primary sugars (maltose, sucrose) were rapidly depleted — particularly under warmer storage — acidogenesis intensified, and acetic acid accumulated (Fig. 2d, S1a). Acetic acid, due to its membrane permeability and higher dissociation constant, imposes greater intracellular stress than lactic acid, which can promote decline of acid-sensitive LAB such as *F. sanfranciscensis* and facilitating the expansion of more acid-tolerant taxa like *A. cerevisiae* (López et al., 2021; Minervini et al., 2014). In parallel, metabolites produced by LAB and yeasts — such as lactic acid, ethanol, and amino acids — likely fostered cross-feeding by AABs, in line with established models of microbial succession involving AABs in other fermentation ecosystems (Blasche et al., 2021).

In parallel, untargeted FIA-MS metabolomics revealed a temperature- and time-dependent enrichment of nucleotides and stress-associated lipid classes (e.g., glycerophospholipids), indicative of nucleic acid degradation, membrane remodeling, and microbial turnover (Fig. 5b–d). The accumulation of select amino acids likely reflects both proteolytic activity and metabolite recycling under nutrient-depleted, acidified conditions (Gänzle & Follador, 2012). Notably, glucose and fructose re-emerged at later timepoints (Fig. S3c), possibly released via endogenous carbohydrate breakdown or also via microbial lysis. These catabolic byproducts may have enabled the metabolic expansion of *A. cerevisiae*, which not only exhibits high stress tolerance but, as observed in kefir systems (Blasche et al., 2021) may also represent a late-successional taxon in the sourdough ecosystem that capitalizes on nutrient landscapes shaped by earlier fermenters.

Despite pronounced metabolic and bacterial shifts, fungal community structure remained remarkably stable across conditions (Figs. 4, 5), highlighting a functional decoupling between taxonomic stability and functional activity. This supports the notion that community composition does not always predict ecosystem function — a phenomenon previously reported in, e.g., marine microbiomes (Louca et al., 2018).

However, this decoupling was asymmetric: in bacterial communities, structural and functional profiles became more tightly coupled under warm, nutrient-limited conditions. Procrustes analyses revealed the strongest alignment between bacterial composition and metabolome profiles at 17 °C and 30 °C (M² ≈ 0.44–0.59; Fig. 5a), suggesting that under increased stress, microbial structure becomes a better predictor of function. This tighter coupling may arise from the erosion of functional redundancy under stress conditions.

Simultaneously, starvation stress exposure increased bacterial beta-dispersion (PERMDISP p < 0.001; Fig. 4), consistent with greater stochasticity and adaptive bottlenecks (Shade et al., 2012; Stegen et al., 2012). These shifts imply that while function becomes more predictable from community structure under stress, the structure itself becomes more variable between replicates. Temperature effects explained progressively more variance in Bray-Curtis dissimilarity (nested PERMANOVA R²: 30 °C > 17 °C > 4 °C), underscoring the dominant role of environmental filtering in shaping microbial ecology during storage.

Multi-omics network analyses revealed profound rewiring of ecological interaction topologies in response to storage temperature. At 4 °C, networks exhibited a modular architecture with *F. sanfranciscensis* maintaining high abundance but lower connectivity, consistent with niche partitioning and community stability under cold, sugar-rich conditions (Fig. 6c,f; S9, S10). In contrast, elevated temperatures led to a collapse of modularity and emergence of centralized structures, where *A. cerevisiae* assumed a hub-like role with high betweenness and eigenvector centrality. This transition likely illustrates a stress-induced reorganization from distributed, niche-driven coexistence to centralized ecological dependence, with *A. cerevisiae* functioning as an interaction keystone — likely mediating metabolite fluxes or coordinating inter-kingdom dynamics, supporting its previously suggested metabolic importance when studied in synthetic sourdough co-cultures (Rappaport et al., 2024).

These shifts were uncovered through integrated, multi-layer network models that capture both positive and negative associations among bacteria, fungi, and metabolites. Unlike single-layer approaches, these frameworks reveal both abundance correlations and structural embedding — linking taxa to metabolite turnover and ecological roles. Notably, *A. cerevisiae*’s increased centrality did not merely reflect growth, but highlighted its functional anchoring as a metabolic and ecological mediator within the network at elevated temperatures. This underscores how community dominance under, e.g. stress conditions, can emerge not only from abundance, but from a taxon’s network position and embeddedness. Such multi-domain integration aligns with recent advances in gut microbiome research (Akiyama et al., 2024), and offers a powerful framework for investigating resilience and stability also in fermented food ecosystems.

Building on this systems-level view, we next explored how predictive signatures of storage conditions are embedded within multi-omics profiles. Our supervised learning approach revealed that metabolite features were disproportionately sensitive to storage conditions: 24 of the top 25 predictors of storage temperature and duration were derived from the FIA-MS metabolome (Fig. 7b). This underscores the heightened vulnerability of functional layers to post-collection drift. Metabolomic data thus not only reflect system function but also encode metadata signatures of handling history and post-collection storage bias. Such sensitivity, while a source of distortion, also provides an opportunity — these molecular signatures could serve as proxies for sample integrity or environmental exposure.

These findings carry critical implications for microbiome studies relying on decentralized sampling and transport. Even brief exposure (1–2 days) to elevated temperatures triggered measurable changes in metabolite composition, bacterial evenness, and viability, while prolonged exposure (from 5–6 days onwards) induced divergence in community structure and ecological networks (Fig. 2–6). Notably, microbial succession lagged behind metabolome remodeling, underscoring the heightened sensitivity of the metabolome to handling conditions — and its potential utility as an early marker of post-collection perturbation.

Such storage-induced biases can distort ordination and clustering, mislead feature selection, and undermine the accuracy of downstream machine learning models — especially when unrecognized. Our findings echo concerns raised in other microbiome contexts, including human gut studies, where technical artifacts have been shown to dominate biological signals if uncorrected (Amir et al., 2017; McLaren et al., 2019; Sinha et al., 2017). These observations emphasize the urgent need for rigorous metadata tracking, awareness of post-collection shifts, and the development of standardized correction workflows — particularly in large-scale, decentralized food microbiome efforts.

In conclusion, our study combines high-resolution storage tracking with machine learning–driven modeling in a controlled sourdough ecosystem to illuminate how microbial and metabolic profiles shift during post-collection storage. Through integrative, multi-omics analysis, we present a scalable, generalizable framework for quantifying and modeling collection-induced bias in microbiome research, and sourdough proved again its value as a tractable model for studying ecological and functional dynamics under environmental stress. Critically, the strong predictive signals encoded in storage-altered multi-omics profiles offer a new avenue for detecting — and potentially correcting — post-collection drift. This opens the door to more accurate, bias-aware interpretation and comparability of microbiome multi-omics data from decentralized, large-scale studies across diverse systems.

## Materials and methods

### Fermentation Experiments

To simulate transportation time and seasonal temperature variation, sourdough samples were stored at 4 °C, 17 °C, and 30 °C, and sampled at 11 time points over one month (daily for the first 7 days, then on days 10, 14, 21, and 28). A laboratory-prepared wheat sourdough starter (dough yield 200, i.e., 100 grams of organic wheat flour (L170524, Swissmill, CH-8037, Zurich) in 100 mL of autoclaved, filtered water) was upscaled overnight to 8 kg (5 containers, each inoculated with 200 g of the same mother sourdough, 700 g of water and 700 g of flour, incubated at 20 °C, 80% humidity) and divided into 99 aliquots (Gosselin Straight Containers, Fisher Scientific AG, Cat. No. 15458824) of 75 g to cover the three storage temperatures and 11 sampling points. Samples were incubated with closed lids until harvest.

### Sample collection and processing

At each time point, three aliquots per temperature condition were randomly selected for microbiological and chemical analyses. All samples (n = 99), including the baseline sourdough (day 0), were diluted 1:1 in molecular-grade water and stored at −80 °C for subsequent DNA and metabolite extraction. For microbiological analysis, 5 g of sourdough was homogenized with 45 g of 1% vegetable peptone (VWR International, Cat. Nr. OXOIVG0100B) and 0.85% NaCl solution, then serially diluted. Viable lactic acid bacteria (LAB) counts were determined by plating on De Man, Rogosa, and Sharpe 5 (MRS-5) agar (Harth et al., 2016), prepared following their exact recipe containing 0.1 g/L cycloheximide, except exchanging meat extract with Beef Extract 500 (Merck, Cat. Nr. B4888-500G), and peptone with vegetable peptone (VWR International, Cat. Nr. OXOIVG0100B). Viable yeast counts were determined by plating on Yeast Extract Peptone Glucose (YPG) agar supplemented with 0.2 g/L chloramphenicol. Plates were incubated aerobically at 30 °C and 80% relative humidity for 48 h before calculating the number of colony-forming units (CFU) per gram. For chemical analysis, 10 g of sourdough was homogenized with 100 g of non-sterile deionized water. The pH was measured immediately, and total titratable acidity (TTA) was quantified by titrating with 0.1 M NaOH to a pH endpoint of 8.5 (Nielsen, 2010).

### Substrate removal from sourdough samples and DNA extraction

DNA was extracted using the MagMAX™ Microbiome Ultra Nucleic Acid Isolation Kit (Thermo Fisher Scientific) on the KingFisher Apex platform (Thermo Fisher Scientific), following the MagMAX Liquid Buccal protocol (Thermo Fisher Scientific) with minor modifications. Cryotubes containing sourdough aliquots were thawed on ice, randomized, and transferred to KingFisher 96 deep-well plates. ZymoBIOMICS Microbial Community Standard (37.5 μL diluted in 112.5 μL sterile water; Zymo Research, Cat. No. D6300) and sterile H₂O (150 μL) served as positive and negative extraction controls, respectively. To prevent yeast sedimentation, samples were briefly homogenized at 20 Hz using a TissueLyser III (Qiagen). To remove flour particles while retaining larger yeast cells, plates were centrifuged at 100 × g for 1 min. A 150 μL aliquot of the resulting supernatant was transferred to the MagMAX Bead Beating Plate. Bead beating was performed twice for 5 minutes at 30 Hz, and DNA was eluted in 50 μL of elution buffer. DNA concentrations were measured using the Qubit™ dsDNA Quantification Assay Kit (Thermo Fisher Scientific, Cat. No. Q32854), following the manufacturer’s instructions. DNA was stored at −20 °C until further processing.

### Marker gene amplicon library preparation and sequencing

Amplicon libraries for bacterial (16S rRNA gene) and fungal (ITS) profiling were prepared using the HighALPS ultra-high-throughput protocol with unique dual indices (UDI) (Flörl et al., 2024). Bacterial communities were amplified targeting the V4 region using UDI-linked primers 515F (5′- GTGYCAGCMGCCGCGGTAA-3′) and 806R (5′-GGACTACNVGGGTWTCTAAT-3′) (Apprill et al., 2015; Parada et al., 2016). Reactions (25 μL) contained 5 μL of template DNA, 12.5 μL 2× KAPA HiFi HotStart ReadyMix (Roche, Cat. No. 07958935001), and 0.3 μM primers. PCR conditions were: 95 °C for 5 min; 35 cycles of 98 °C for 20 s, 55 °C for 15 s, and 72 °C for 25 s; followed by a final extension at 72 °C for 4 min.

Fungal communities were profiled via nested PCR targeting the ITS1 region using BITS (5′- ACCTGCGGARGGATCA-3′) and B58S3 (5′-GAGATCCRTTGYTRAAAGTT-3′) primers (Bokulich & Mills, 2013). The first round used non-indexed primers (25 μL reactions with 2 μL DNA template), followed by barcoding PCR using UDI-tagged primers with 1 μL of the first reaction as input. All primers were ordered from Microsynth AG, Switzerland. Cycling conditions were: 95 °C for 5 min; 35 cycles of 98 °C for 20 s, 49 °C for 15 s, and 72 °C for 20 s; with a final extension at 72 °C for 4 min. Barcoding PCR consisted of 10 cycles under the same thermal profile.

Amplicons were purified using Agencourt AMPure XP magnetic beads (0.7× ratio; Beckman Coulter, Cat. No. A63882) on the KingFisher Apex platform. DNA concentrations were quantified in duplicate using the Qubit™ dsDNA High Sensitivity Assay (Thermo Fisher Scientific, Cat. No. Q32854) on a Tecan Spark microplate reader. 16S rRNA and ITS amplicons were pooled separately at equimolar concentrations using a liquid handling platform (Brand GmbH), and quality was assessed via high-sensitivity TapeStation (Agilent Technologies, High Sensitivity D1000 DNA ScreenTape assays, Cat. No. 5067-5584). Pooled libraries were combined and sequenced with 300 bp paired-end reads on the Illumina NextSeq 2000 (600-cycle kit), including a 20% PhiX spike-in, at the Functional Genomics Center Zürich.

### Quantitative PCR

Quantitative PCR (qPCR) was used to determine total, viable and non-viable, bacterial (LAB and non-LAB) abundance across all storage conditions. Each 25 μL reaction contained 1 μL of template DNA, 12.5 μL of 2× KAPA HiFi HotStart ReadyMix (Roche, Cat. No. 07958935001), 1 μL of 20× EvaGreen Dye (Brunschwig, Cat. No. BIO31000), and 0.3 μM of non-barcoded 515F/806R primers. Thermal cycling was performed on a LightCycler 480 (Roche) with the following conditions: 95 °C for 3 min; 40 cycles of 98 °C for 20 s, 55 °C for 15 s, and 72 °C for 15 s; followed by melt curve analysis (95 °C for 30 s, 60 °C for 15 s, ramping at 1 °C/min). Absolute DNA concentrations (ng/μL) were determined using a standard curve generated with the ZymoBIOMICS Microbial Community DNA Standard (Zymo Research, Cat. No. D6305), and applied to cycle threshold (Ct) values from sourdough samples. Bacterial cell counts were estimated assuming an average of five 16S rRNA gene copies per cell and 1.58 × 10⁹ 16S rRNA gene copies per ng of DNA (Table S1).

### HPLC sample preparation and measurement

Pre-diluted sourdough samples stored at −80 °C were thawed on ice for 30–45 min. A 200 mg aliquot was diluted 1:10 in LC-MS grade water (Merck Supelco), homogenized in a Thermomix at 1500 rpm and 10 °C for 10 min, and centrifuged at 14,000 × g and 4 °C for 15 min. The resulting supernatant was filtered through a 0.45 μm polyvinylidene difluoride (PVDF) membrane (BGB Analytik AG) to remove particulates. For organic acid and ethanol analysis, 200 μL of filtered supernatant was transferred into glass HPLC vials with inserts. For sugar analysis, 100 μL of supernatant was further diluted 1:10 with Milli-Q water (MilliporeSigma) in plastic vials. Samples were stored at 4 °C for up to 48 h or at −20 °C for longer-term storage.

### Organic acid and ethanol detection

Organic acids (succinic acid, lactic acid, acetic acid) and ethanol were quantified using an Agilent 1200 Series HPLC system equipped with an Aminex HPX-87H column (Bio-Rad). The column was maintained at 40 °C with a sample tray temperature of 10 °C and an injection volume of 10 μL. Separation was achieved under isocratic conditions using 5 mM sulfuric acid (H₂SO₄; Merck Titrisol) at a flow rate of 0.4 mL/min over 40 min. Ethanol was detected via refractive index (RI) detection at 40 °C, while organic acids were quantified using a diode array detector at 210 nm.

### Carbohydrate detection

Carbohydrates (mannitol, sucrose, glucose, fructose, maltose) were analyzed using a Dionex ICS-5000+ system (Thermo Scientific) equipped with a Dionex CarboPac PA200 IC column maintained at 25 °C. Samples were held at 10 °C and injected at 10 μL. Separation was performed at a flow rate of 0.25 mL/min using a three-solvent system: (1) 225 mM sodium hydroxide (NaOH), (2) 1250 mM sodium acetate with 100 mM NaOH, and (3) Milli-Q water. A defined gradient program (Table S2) was applied, and carbohydrates were detected via pulsed amperometric detection. Total runtime per sample was 45 min.

### FIA-MS untargeted metabolomics

All steps were performed on ice using ice-cold solvents unless otherwise specified. For metabolite fingerprinting via flow injection analysis-mass spectrometry (FIA-MS), 100 μL of water extract (from the HPLC organic acid analysis; see above) was diluted with 900 μL LC-MS grade methanol (Fisher Chemical), vortexed briefly, and centrifuged at 4,500 × g for 30 min at 5 °C. The supernatant was further diluted 1:10 in 50% methanol, mixed, and 150 μL was transferred to a 96-well plate, heat-sealed with foil (Thermo Fisher Scientific), and analyzed within 24 h. FIA-MS was conducted using an Acquity I-Class UPLC system coupled to a Xevo G2-XS qToF mass spectrometer (Waters), following a modified protocol from Fuhrer et al. (2011). Two microliters of extract were injected in 60% methanol containing 0.05% ammonium hydroxide (NH₄OH, Honeywell) and 2 mM ammonium fluoride (NH₄F; Sigma-Aldrich) as carrier solvent, at 0.2 mL/min for 1 min. Compounds were ionized using electrospray ionization in negative sensitivity mode with the following source parameters: capillary voltage 2.2 kV, cone voltage 40 V, source offset 80 V, source temperature 120 °C, desolvation gas at 250 °C and 850 L/h, and cone gas at 150 L/h. Mass spectra were acquired in extended dynamic range from m/z 50–1200 at a scan rate of 0.7 s. For online mass correction, 200 ng/μL Leucine-Enkephalin ([M–H]⁻ 556.2771; Waters) in 50:50:0.1 acetonitrile:water:formic acid was injected every 19 s (scan time 0.3 s, capillary voltage 2.2 kV).

Raw data were converted to mzML format using MSConvert v3.0 (ProteoWizard), and processed in MZMine 4.1.0 (Schmid et al., 2023). Mass detection used centroid mode with a noise level of 1000. Chromatogram building used the ADAP algorithm (min scans: 4, min group intensity: 2000, min absolute height: 2000, m/z tolerance: 0.005 or 10 ppm). Chromatograms were smoothed with a Savitzky–Golay filter (RT window: 5). Carbon-13 deisotoping applied intra-sample tolerances of 0.002 m/z or 3.5 ppm and 1 min RT. Feature alignment used the Join Aligner (m/z tolerance: 0.005 or 15 ppm; m/z weight: 80; RT tolerance: 1 min; RT and mobility weights: 1). Gap filling used an intensity tolerance of 0.2, m/z tolerance of 0.005 or 15 ppm, and a minimum of 4 scans. Blank subtraction removed features with intensity <5x that of corresponding blanks. Final exported peak heights were used for statistical analysis.

Ions were annotated by exact mass (m/z) matching (±0.002 Da) against the KEGG metabolite (Kanehisa et al., 2016) and HMDB databases (Wishart et al., 2018), and subsequently classified using the HMDB metabolite taxonomy. Annotated features were then grouped into compound classes or subclasses relevant to sourdough fermentation (e.g., amino acids, organic acids, carbohydrates) for visualization. These groupings were used to calculate and plot the relative proportions of annotated metabolites within each temporal response cluster and temperature condition (Fig. 4).

### Bioinformatic marker gene amplicon processing

#### Fungal ITS amplicons

Raw paired-end internal transcribed spacer (ITS) sequences were processed using QIIME 2 version 2024.10 (Bolyen et al., 2019). Primers and adapters were trimmed with the cutadapt trim-paired plugin (Martin, 2011). Denoising was performed via dada2 denoise-single plugin (Callahan et al., 2016) with no truncation (--p-trunc-len 0), a maximum expected error of 4.0, and a minimum parent abundance fold-difference of 4.0, resulting in 96–98% non-chimeric reads. Reads shorter than 50 nt were filtered out, retaining 2,453,636 sequences across 33 samples per temperature group (4 °C, 17 °C, 30 °C), one day 0 mother control, and excluding one contaminated replicate at day 7. Taxonomy was assigned with the classify-sklearn action of the q2-feature-classifier plugin (Bokulich et al., 2018) against the customized UNITE v10.99 reference database (Abarenkov et al., 2024) curated via RESCRIPt (Robeson et al., 2021), with a confidence threshold of 0 to retain low-confidence fungal reads. Reads assigned to non-fungal sequences, low-abundance features (<10 reads/sample), fruiting body–associated taxa, unclassified phyla, and extremely rare class-level taxa were removed, accounting for 1.4% of total reads and resulting in 2,419,268 reads. Operational taxonomic units (OTUs) were generated by mapping reads at 90% similarity using the vsearch cluster-features-closed-reference plugin (Rognes et al., 2016), against the same UNITE v10.99 database, excluding singletons. Unmatched and poorly classified reads were discarded, yielding a final OTU table of 2,411,351 reads—a 0.33% reduction from the ASV set. Both ASV and OTU tables were rarefied to 3,500 reads per sample, resulting in a final dataset of 95 samples (Fig. S14a,b).

#### Bacterial 16S rRNAv4 amplicons

Paired-end 16S rRNAv4 gene sequences were processed using the dada2 denoise-paired plugin in QIIME 2. Reads were truncated at 150 bp in both directions to remove low-quality regions, primers, and adapters. Denoising used dada2 (via the q2-dada2 plugin) with a maximum expected error threshold of 2.0 and a minimum fold-parent-over-abundance threshold of 4.0, yielding 84– 86% non-chimeric reads and a total of 4,019,885 denoised, merged sequences. One sample (day 28 at 4 °C) failed, resulting in 32 samples for 4 °C, 33 for 17 °C, 33 for 30 °C, and 1 mother sourdough control. All retained reads were ≥50 bp, so no additional length filtering was required. Taxonomic classification was performed using the classify-sklearn plugin trained on a custom SILVA 138.2 SSU NR99 reference database (Quast et al., 2013; Yilmaz et al., 2014), restricted to EMP 515f/806r amplicon regions and prepared using RESCRIPt. A confidence threshold of 0 was used to maximize bacterial read retention and to only exclude non-bacterial sequences (e.g., mitochondria, chloroplasts, archaea, eukaryotes), which accounted for 13.46% of reads. An additional 492 ASVs were removed via contaminant filtering (Davis et al., 2018) (decontam-combined action of the q2-quality-control plugin) and removal of rare features (<10 reads in any sample), resulting in 3,478,383 high-quality reads. Reads were additionally clustered into OTUs at 99% identity using the vsearch cluster-features-closed-reference plugin against SILVA 138.2, yielding a final OTU table with 3,478,356 reads (only 27 fewer than the ASV table due to unmatched sequences). Both ASV and OTU tables were subsequently rarefied to 380 reads/sample, resulting in a final dataset of 93 samples (Fig. S14c,d).

### Statistical analysis

#### Microbial Diversity analyses

Diversity analyses, including calculations of alpha-diversity (richness, evenness and Shannon entropy) and beta-diversity (Jaccard distance and Bray–Curtis dissimilarity), were made using the q2-diversity and q2-kmerizer (Bokulich, 2025) plugins in QIIME 2. Beta diversity estimates were calculated based on ASV composition as well as the constituent k-mers in these ASVs, and OTU features.

#### FIA-MS features preprocessing and clustering

FIA-MS features were filtered with a prevalence threshold of 3 (triplicates) and missing values were imputed as zeros. Feature tables were processed using z-score normalization, log transformation, combined log-z transformation, or total sum scaling (TSS). To capture dynamic metabolite responses to storage, one-way ANOVA (α = 0.05) was performed across time points within each temperature group on z-score normalized data. Metabolites showing significant temporal variation in at least one temperature condition were retained, and non-responsive features were excluded prior to clustering. Filtered, z-score normalized metabolite features were clustered using k-means in scikit-learn (MacQueen, 1967; Pedregosa et al., 2011). Optimal cluster number was determined via silhouette score, elbow method (WCSS), and Gap statistic. Final clusters were defined with the best-supported k (k = 3) across all evaluation metrics. For each cluster, temporal dynamics were visualized by plotting z-score trajectories per temperature. Kruskal-Wallis tests were performed per cluster and day to assess temperature-driven differences, with Benjamini-Hochberg correction applied to control for false discovery rate.

#### Two-dimensional feature space statistics

Microbial beta-diversity feature spaces were visualized by using principal coordinate analysis (PCoA) and FIA-features by using Principal Component Analysis (PCA) to reduce the dimensionality. To aid interpretation of sample separation in beta-diversity PCoAs and FIA PCAs, top correlated features and metadata variables were overlaid in feature space. Pearson correlations between each variable and the first two PCoA respectively PCA axes were used to determine direction and length. Top contributors were visualized as scaled vectors, highlighting features and metadata most associated with ordination structure. To evaluate the contributions of experimental factors and their nested or interacting effects on community and metabolomic dissimilarity structures, we applied the QIIME 2 diversity adonis plugin for global PERMANOVA modeling (Fig. S7a,d,g) and a custom two-level nested PERMANOVA implementation in scikit-bio (Fig. S7b,c,e,f,h,i) (Anderson, 2001; Oksanen et al., 2001; Rideout et al., 2023). For the adapted nesting, global variance was first assessed for primary factors (e.g., temperature or day), followed by separate within-group PERMANOVA tests on secondary factors (e.g., day nested within each temperature, or temperature nested within each day). p-values were corrected for multiple testing using the Benjamini–Hochberg false discovery rate (FDR) procedure.

#### Day-wise comparison with post hoc grouping

To assess temperature effects across time, one-way ANOVA was applied at each time point for continuous variables including alpha diversity (amplicon-based and FIA-MS), CFUs, pH, TTA, and HPLC-derived metabolite concentrations (excluding control samples). Post hoc comparisons were performed using Tukey’s Honest Significant Difference (HSD) test to identify temperature-specific differences per day. Prior to conducting ANOVAs, homogeneity of variance was evaluated using Levene’s test across temperatures for each time point and variable (Fig. S15). As no comparisons showed significant heterogeneity (p < 0.05), ANOVA assumptions were considered met despite small replicate numbers.

#### Differential abundance analysis

Microbial compositional shifts were tested using ANCOM-BC2 (Lin & Peddada, 2024) in R v4.5.0. Amplicon data were rescaled to pseudo-counts and analyzed using a phyloseq-based workflow, excluding control (20 °C) samples. The model included day and temperature as fixed effects, and both pairwise and global tests were performed with FDR correction. Structural zeros and sampling fraction variation were accounted for. Taxa showing log-fold changes with a certain threshold (|logFC| > 0.75 for 16S, |logFC| > 0.1 for ITS) were classified as enriched (numerator) or outcompeted (denominator). Sample-wise log-ratios were computed as the log geometric mean of enriched taxa over depleted taxa at given temperatures and time points.

#### Integrations of metabolomics and amplicon data

To assess the concordance between metabolic and microbial community structures, Bray-Curtis dissimilarity matrices were computed on 16S OTU feature tables and subsequent PCoA, whereas FIA-MS (either z-score or TSS normalized) was either processed via Bray-Curtis and PCoA or direct PCA. Subsequently, Procrustes analysis was used to align ordination spaces between microbiome and metabolome data. To evaluate statistical significance, permutation tests (n = 1,000) were conducted by randomizing sample labels in one dataset and recomputing Procrustes disparities, yielding empirical p-values. Disparities and associated p-values were computed for global comparisons, as well as stratified by day and temperature in isolation or stratification, to identify temporal or condition-specific shifts in metabolome–microbiome concordance.

#### Multi-omics co-occurrence analysis and network inference

To investigate intra- and inter-kingdom associations in the sourdough microbiome and metabolome across storage temperatures, we performed two complementary network inference approaches based on centered log-ratio (CLR)–transformed data. In the first approach, microbial OTU tables (16S and ITS) and metabolomics data were CLR-transformed (after multiplicative replacement; delta=1e-8; with TSS normalization for metabolite data) and pairwise associations were calculated using a novel abundance-weighted co-occurrence score:

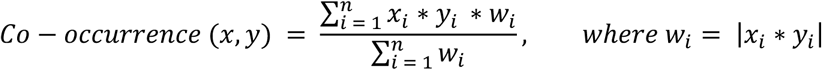

This formula, inspired by compositional-aware correlation frameworks such as SparCC (Friedman & Alm, 2012) and CoNet (Faust & Raes, 2016), enhances traditional co-occurrence metrics by explicitly upweighting high-abundance, co-varying features while suppressing noise from low-abundance artifacts. Unlike SparCC, which infers correlations via iterative pseudo-count-based covariance estimation, or CoNet, which combines multiple similarity metrics with heuristic edge selection, our score directly integrates abundance information into the association weight, offering greater sensitivity to biologically meaningful, high-confidence co-occurrence in complex multi-omics datasets. Associations with ∣Score(x,y)∣ ≥ 0.2 were retained for downstream network construction. This approach supports robust detection of ecologically relevant interactions across kingdoms and omics layers, while maintaining interpretability and scalability for large datasets.

In parallel, we applied a sparse inverse covariance estimation approach to capture conditional dependencies and partial correlations among features. CLR-transformed feature matrices were z-score standardized and concatenated, followed by model fitting using the Graphical Lasso with cross-validation in scikit-learn (Friedman et al., 2008; Pedregosa et al., 2011). Non-zero entries in the resulting precision matrix defined network edges, representing regularized partial correlations. Feature identity (bacteria, fungi, metabolite) was used to classify intra- and inter-domain interactions. The resulting co-occurrence matrices were visualized as undirected networks in NetworkX (v2.8.8) (Hagberg et al., 2008), with node attributes encoding biological category (bacteria, fungi, metabolite), shape, and average abundance (used for node size scaling). Network layout was performed using Graphviz’s “neato” force-directed algorithm (Gansner & North, 2000). Edge thickness and spring length were scaled by co-occurrence strength, and edge color represented co-occurrence polarity. Topological metrics - degree centrality, betweenness centrality, closeness centrality, and eigenvector centrality - were calculated to identify key taxa and metabolites in each network. Networks and centrality metrics were generated independently for each storage temperature.

#### Supervised classification of storage conditions using multi-omics features

To predict sample storage temperature (4 °C, 17 °C, 30 °C), a supervised classifier was trained on combined microbial (16S OTU) and metabolomic (FIA-MS) feature matrices as well as HPLC, CFU, pH, TTA and DNA concentration (extractions) data. A nested pipeline was implemented using a random forest classifier (Breiman, 2001) (100 estimators) preceded by feature selection, which retained features with importances above the median threshold. Model evaluation was performed using stratified 5-fold cross-validation to preserve class balance per fold. This approach allowed robust performance estimation while accounting for both model fitting and feature selection within each training fold.

All statistical and computational analyses were performed in Python (v3.10.14), unless stated otherwise. Data preprocessing used pandas (The pandas development team, 2024) and numpy (Harris et al., 2020). Dimensionality reduction and building and evaluating random forest models was performed with scikit-learn (Pedregosa et al., 2011). Random forest models were built and evaluated using scikit-learn. Statistical modeling (e.g., ANOVA) used statsmodels (Seabold & Perktold, 2010). Visualizations were generated with Seaborn (Waskom, 2021), Matplotlib (The Matplotlib Development Team, 2024) and NetworkX (Hagberg et al., 2008).

## Acknowledgements

The authors acknowledge financial support from the project HealthFerm, which is funded by the European Union under the Horizon Europe grant agreement No. 101060247 and by the Swiss State Secretariat for Education, Research and Innovation (SERI) under contract No. 22.00210. Views and opinions expressed are however those of the author(s) only and do not necessarily reflect those of the European Union nor European Research Executive Agency (REA). Neither the European Union nor REA can be held responsible for them.

The authors thank Luisa Ferreira for support in data collection and the Genomic Diversity Center of ETH Zürich for their support with amplicon library preparation. The microbiome amplicon sequencing was performed at the Functional Genomics Center Zurich of University of Zurich and ETH Zurich.

## Author contributions

Annina R. Meyer, Conceptualization, Formal analysis, Investigation, Visualization, Methodology, Writing - original draft, Writing - review and editing; Jan P. Tan, Formal analysis, Investigation, Methodology, Writing - review and editing; Mihnea P. Mihaila, Michelle Neugebauer, Investigation, Writing - review and editing; Laura Nyström, Resources, Supervision, Funding acquisition, Writing - review and editing; Nicholas A. Bokulich, Conceptualization, Resources, Supervision, Funding acquisition, Writing - original draft, Writing - review and editing.

## Data availability

All sequence data have been deposited on EBI-ENA under accession number PRJEB94514 (16S) and PRJEB94515 (ITS). Source data (metadata) along with processed HPLC and FIA-MS data have been deposited together with code notebooks in github https://github.com/anninameyer/shipped-and-shifted.

## References

Akiyama, S., Nishijima, S., Kojima, Y., Kimura, M., Ohsugi, M., Ueki, K., Mizokami, M., Hattori, M., Tsuchiya, K., Uemura, N., Kawai, T., Bork, P., & Nagata, N. (2024). Multi-biome analysis identifies distinct gut microbial signatures and their crosstalk in ulcerative colitis and Crohn’s disease. Nature Communications, 15(1), 10291. 10.1038/s41467-024-54797-8

Amir, A., McDonald, D., Navas-Molina, J. A., Debelius, J., Morton, J. T., Hyde, E., Robbins-Pianka, A., & Knight, R. (2017). Correcting for Microbial Blooms in Fecal Samples during Room-Temperature Shipping. mSystems, 2(2), 10.1128/msystems.00199-16.

Baig, M. A., Turner, M. S., Liu, S.-Q., Al-Nabulsi, A. A., Shah, N. P., & Ayyash, M. M. (2021). Potential Probiotic Pediococcus pentosaceus M41 Modulates Its Proteome Differentially for Tolerances Against Heat, Cold, Acid, and Bile Stresses. Frontiers in Microbiology, 12. 10.3389/fmicb.2021.731410

Banerjee, S., Walder, F., Büchi, L., Meyer, M., Held, A. Y., Gattinger, A., Keller, T., Charles, R., & van der Heijden, M. G. A. (2019). Agricultural intensification reduces microbial network complexity and the abundance of keystone taxa in roots. The ISME Journal, 13(7), 1722–1736. 10.1038/s41396-019-0383-2

Bassis, C. M., Moore, N. M., Lolans, K., Seekatz, A. M., Weinstein, R. A., Young, V. B., Hayden, M. K., & for the CDC Prevention Epicenters Program. (2017). Comparison of stool versus rectal swab samples and storage conditions on bacterial community profiles. BMC Microbiology, 17(1), 78. 10.1186/s12866-017-0983-9

Berg, G., Rybakova, D., Fischer, D., Cernava, T., Vergès, M.-C. C., Charles, T., Chen, X., Cocolin, L., Eversole, K., Corral, G. H., Kazou, M., Kinkel, L., Lange, L., Lima, N., Loy, A., Macklin, J. A., Maguin, E., Mauchline, T., McClure, R., … Schloter, M. (2020). Microbiome definition re-visited: Old concepts and new challenges. Microbiome, 8(1), 103. 10.1186/s40168-020-00875-0

Blasche, S., Kim, Y., Mars, R. A. T., Machado, D., Maansson, M., Kafkia, E., Milanese, A., Zeller, G., Teusink, B., Nielsen, J., Benes, V., Neves, R., Sauer, U., & Patil, K. R. (2021). Metabolic cooperation and spatiotemporal niche partitioning in a kefir microbial community. Nature Microbiology, 6(2), 196–208. 10.1038/s41564-020-00816-5

Bokulich, N. A., Maldonado, J., Kang, D.-W., Krajmalnik-Brown, R., & Caporaso, J. G. (2019). Rapidly Processed Stool Swabs Approximate Stool Microbiota Profiles. mSphere, 4(2), 10.1128/msphere.00208-19.

Burman, E., & Bengtsson-Palme, J. (2021). Microbial Community Interactions Are Sensitive to Small Changes in Temperature. Frontiers in Microbiology, 12. 10.3389/fmicb.2021.672910

Cabello-Olmo, M., Oneca, M., Torre, P., Díaz, J. V., Encio, I. J., Barajas, M., & Araña, M. (2020). Influence of Storage Temperature and Packaging on Bacteria and Yeast Viability in a Plant-Based Fermented Food. Foods, 9(3), 302. 10.3390/foods9030302

Corsetti, A., & Settanni, L. (2007). Lactobacilli in sourdough fermentation. Food Research International, 40(5), 539–558. Scopus. 10.1016/j.foodres.2006.11.001

De Filippis, F., Genovese, A., Ferranti, P., Gilbert, J. A., & Ercolini, D. (2016). Metatranscriptomics reveals temperature-driven functional changes in microbiome impacting cheese maturation rate. Scientific Reports, 6(1), 21871. 10.1038/srep21871

De Vuyst, L., Van Kerrebroeck, S., & Leroy, F. (2017). Chapter Two—Microbial Ecology and Process Technology of Sourdough Fermentation. In S. Sariaslani & G. M. Gadd (Eds.), Advances in Applied Microbiology (Vol. 100, pp. 49–160). Academic Press. 10.1016/bs.aambs.2017.02.003

Ercolini, D., Pontonio, E., De Filippis, F., Minervini, F., La Storia, A., Gobbetti, M., & Di Cagno, R. (2013). Microbial Ecology Dynamics during Rye and Wheat Sourdough Preparation. Applied and Environmental Microbiology, 79(24), 7827–7836. 10.1128/AEM.02955-13

Gänzle, M., & Follador, R. (2012). Metabolism of Oligosaccharides and Starch in Lactobacilli: A Review. Frontiers in Microbiology, 3. 10.3389/fmicb.2012.00340

Gilbert, J. A., Blaser, M. J., Caporaso, J. G., Jansson, J. K., Lynch, S. V., & Knight, R. (2018). Current understanding of the human microbiome. Nature Medicine, 24(4), 392–400. 10.1038/nm.4517

Gomes, R. J., Borges, M. de F., Rosa, M. de F., Castro-Gómez, R. J. H., & Spinosa, W. A. (2018). Acetic Acid Bacteria in the Food Industry: Systematics, Characteristics and Applications. Food Technology and Biotechnology, 56(2), 139–151. 10.17113/ftb.56.02.18.5593

Han, N. R., Yu, S., Byun, J. A., Yun, E. J., Cheon, S., Song, S., Shim, S., Choi, I.-G., Lee, S.-H., & Kim, K. H. (2024). Evolution-aided improvement of the acid tolerance of *Levilactobacillus brevis* and its application in sourdough fermentation. Food Research International, 190, 114584. 10.1016/j.foodres.2024.114584

Kilstrup, M., Hammer, K., Ruhdal Jensen, P., & Martinussen, J. (2005). Nucleotide metabolism and its control in lactic acid bacteria. FEMS Microbiology Reviews, 29(3), 555–590. 10.1016/j.fmrre.2005.04.006

Kim, E., Yang, S.-M., & Kim, H.-Y. (2021). Analysis of Cultivable Microbial Community during Kimchi Fermentation Using MALDI-TOF MS. Foods, 10(5), Article 5. 10.3390/foods10051068

Kim, J. Y., Park, S.-E., Kim, E.-J., Seo, S.-H., Whon, T. W., Cho, K.-M., Kwon, S. J., Roh, S. W., & Son, H.-S. (2022). Long-term population dynamics of viable microbes in a closed ecosystem of fermented vegetables. Food Research International, 154, 111044. 10.1016/j.foodres.2022.111044

Landis, E. A., Oliverio, A. M., McKenney, E. A., Nichols, L. M., Kfoury, N., Biango-Daniels, M., Shell, L. K., Madden, A. A., Shapiro, L., Sakunala, S., Drake, K., Robbat, A., Booker, M., Dunn, R. R., Fierer, N., & Wolfe, B. E. (2021). The diversity and function of sourdough starter microbiomes. eLife, 10, e61644. 10.7554/eLife.61644

Lim, J.-Y., Choi, Y.-J., Choi, J.-Y., Yang, J.-H., Chung, Y. B., Park, S.-H., Min, S. G., & Lee, M.-A. (2024). Microbial Dynamics and Metabolite Profiles in Different Types of Salted Seafood (Jeotgal) During Fermentation. ACS Omega, 9(33), 35798–35808. 10.1021/acsomega.4c04410

López, P. C., Peng, C., Arneborg, N., Junicke, H., & Gernaey, K. V. (2021). Analysis of the response of the cell membrane of Saccharomyces cerevisiae during the detoxification of common lignocellulosic inhibitors. Scientific Reports, 11(1), 6853. 10.1038/s41598-021-86135-z

Louca, S., Polz, M. F., Mazel, F., Albright, M. B. N., Huber, J. A., O’Connor, M. I., Ackermann, M., Hahn, A. S., Srivastava, D. S., Crowe, S. A., Doebeli, M., & Parfrey, L. W. (2018). Function and functional redundancy in microbial systems. Nature Ecology & Evolution, 2(6), 936–943. 10.1038/s41559-018-0519-1

Louw, N. L., Lele, K., Ye, R., Edwards, C. B., & Wolfe, B. E. (2023). Microbiome Assembly in Fermented Foods. Annual Review of Microbiology, 77(1), 381–402. 10.1146/annurev-micro-032521-041956

Marco, M. L., Heeney, D., Binda, S., Cifelli, C. J., Cotter, P. D., Foligné, B., Gänzle, M., Kort, R., Pasin, G., Pihlanto, A., Smid, E. J., & Hutkins, R. (2017). Health benefits of fermented foods: Microbiota and beyond. Current Opinion in Biotechnology, 44, 94–102. 10.1016/j.copbio.2016.11.010

Marsland, R. the 3rd, Cui, W., Goldford, J., Sanchez, A., Korolev, K., & Mehta, P. (2019). Available energy fluxes drive a transition in the diversity, stability, and functional structure of microbial communities. PLOS Computational Biology, 15(2), e1006793. 10.1371/journal.pcbi.1006793

Martins, I. E., Shittu, T. A., Onabanjo, O. O., Adesina, A. D., Soares, A. G., Okolie, P. I., Kupoluyi, A. O., Ojo, O. A., & Obadina, A. O. (2021). Effect of packaging materials and storage conditions on the microbial quality of pearl millet sourdough bread. Journal of Food Science and Technology, 58(1), 52–61. 10.1007/s13197-020-04513-3

McDonald, D., Hyde, E., Debelius, J. W., Morton, J. T., Gonzalez, A., Ackermann, G., Aksenov, A. A., Behsaz, B., Brennan, C., Chen, Y., DeRight Goldasich, L., Dorrestein, P. C., Dunn, R. R., Fahimipour, A. K., Gaffney, J., Gilbert, J. A., Gogul, G., Green, J. L., Hugenholtz, P., … Knight, R. (2018). American Gut: An Open Platform for Citizen Science Microbiome Research. mSystems, 3(3), 10.1128/msystems.00031-18.

McLaren, M. R., Willis, A. D., & Callahan, B. J. (2019). Consistent and correctable bias in metagenomic sequencing experiments. eLife, 8, e46923. 10.7554/eLife.46923

Meyer, A., Gettemans, T., Tan, J. P., Tuccillo, F., Viretto, C., Angelescu, I.-R., Bondt, Y. D., Neugebauer, M., Tlais, A. Z. A., Cavelti, F., Vuyst, L. D., Gobbetti, M., Courtin, C. M., Zamfir, M., Coda, R., Nyström, L., Weckx, S., & Bokulich, N. A. (2025). Rising together: Exploring Sourdough Fermentation Diversity through Co-design in the HealthFerm Citizen Science Initiative (p. 2025.05.23.655785). bioRxiv. 10.1101/2025.05.23.655785

Minervini, F., De Angelis, M., Di Cagno, R., & Gobbetti, M. (2014). Ecological parameters influencing microbial diversity and stability of traditional sourdough. International Journal of Food Microbiology, 171, 136–146. 10.1016/j.ijfoodmicro.2013.11.021

Momo Cabrera, P., Bokulich, N. A., & Zimmermann, P. (2025). Evaluating stool microbiome integrity after domestic freezer storage using whole-metagenome sequencing, genome assembly, and antimicrobial resistance gene analysis. Microbiology Spectrum, 13(3), e02278–24. 10.1128/spectrum.02278-24

Papadimitriou, K., Alegría, Á., Bron, P. A., Angelis, M. de, Gobbetti, M., Kleerebezem, M., Lemos, J. A., Linares, D. M., Ross, P., Stanton, C., Turroni, F., Sinderen, D. van, Varmanen, P., Ventura, M., Zúñiga, M., Tsakalidou, E., & Kok, J. (2016). Stress Physiology of Lactic Acid Bacteria. Microbiology and Molecular Biology Reviews : MMBR, 80(3), 837. 10.1128/MMBR.00076-15

Rappaport, H. B., Senewiratne, N. P. J., Lucas, S. K., Wolfe, B. E., & Oliverio, A. M. (2024). Genomics and synthetic community experiments uncover the key metabolic roles of acetic acid bacteria in sourdough starter microbiomes. mSystems, 0(0), e00537–24. 10.1128/msystems.00537-24

Reese, A. T., Madden, A. A., Joossens, M., Lacaze, G., & Dunn, R. R. (2020). Influences of Ingredients and Bakers on the Bacteria and Fungi in Sourdough Starters and Bread. mSphere, 5(1), e00950–19. 10.1128/mSphere.00950-19

Ripari, V., Gänzle, M. G., & Berardi, E. (2016). Evolution of sourdough microbiota in spontaneous sourdoughs started with different plant materials. International Journal of Food Microbiology, 232, 35–42. 10.1016/j.ijfoodmicro.2016.05.025

Sanmartin C, A. G. (2013). The Kinetics of Fermentations in Sourdough Bread Stored at Different Temperature and Influence on Bread Quality. Journal of Bioprocessing & Biotechniques, 03(03). 10.4172/2155-9821.1000134

Sawant, S. S., Park, H.-Y., Sim, E.-Y., Kim, H.-S., & Choi, H.-S. (2025). Microbial Fermentation in Food: Impact on Functional Properties and Nutritional Enhancement—A Review of Recent Developments. Fermentation, 11(1), Article 1. 10.3390/fermentation11010015

Scofield, V., Jacques, S. M. S., Guimarães, J. R. D., & Farjalla, V. F. (2015). Potential changes in bacterial metabolism associated with increased water temperature and nutrient inputs in tropical humic lagoons. Frontiers in Microbiology, 6. 10.3389/fmicb.2015.00310

Shade, A., Peter, H., Allison, S. D., Baho, D., Berga, M., Buergmann, H., Huber, D. H., Langenheder, S., Lennon, J. T., Martiny, J. B., Matulich, K. L., Schmidt, T. M., & Handelsman, J. (2012). Fundamentals of Microbial Community Resistance and Resilience. Frontiers in Microbiology, 3. 10.3389/fmicb.2012.00417

Silva, I., Alves, M., Malheiro, C., Silva, A. R. R., Loureiro, S., Henriques, I., & González-Alcaraz, M. N. (2022). Short-Term Responses of Soil Microbial Communities to Changes in Air Temperature, Soil Moisture and UV Radiation. Genes, 13(5), Article 5. 10.3390/genes13050850

Sinha, R., Abu-Ali, G., Vogtmann, E., Fodor, A. A., Ren, B., Amir, A., Schwager, E., Crabtree, J., Ma, S., Abnet, C. C., Knight, R., White, O., & Huttenhower, C. (2017). Assessment of variation in microbial community amplicon sequencing by the Microbiome Quality Control (MBQC) project consortium. Nature Biotechnology, 35(11), 1077–1086. 10.1038/nbt.3981

Song, S. J., Amir, A., Metcalf, J. L., Amato, K. R., Xu, Z. Z., Humphrey, G., & Knight, R. (2016). Preservation Methods Differ in Fecal Microbiome Stability, Affecting Suitability for Field Studies. mSystems, 1(3), 10.1128/msystems.00021-16.

Stegen, J. C., Lin, X., Konopka, A. E., & Fredrickson, J. K. (2012). Stochastic and deterministic assembly processes in subsurface microbial communities. The ISME Journal, 6(9), 1653–1664. 10.1038/ismej.2012.22

Tan, G., Hu, M., Li, X., Li, X., Pan, Z., Li, M., Li, L., Wang, Y., & Zheng, Z. (2022). Microbial Community and Metabolite Dynamics During Soy Sauce Koji Making. Frontiers in Microbiology, 13. 10.3389/fmicb.2022.841529

Tedjo, D. I., Jonkers, D. M. A. E., Savelkoul, P. H., Masclee, A. A., Best, N. van, Pierik, M. J., & Penders, J. (2015). The Effect of Sampling and Storage on the Fecal Microbiota Composition in Healthy and Diseased Subjects. PLOS ONE, 10(5), e0126685. 10.1371/journal.pone.0126685

Valentino, V., Magliulo, R., Farsi, D., Cotter, P. D., O’Sullivan, O., Ercolini, D., & De Filippis, F. (2024). Fermented foods, their microbiome and its potential in boosting human health. Microbial Biotechnology, 17(2), e14428. 10.1111/1751-7915.14428

Van Kerrebroeck, S., Maes, D., & De Vuyst, L. (2017). Sourdoughs as a function of their species diversity and process conditions, a meta-analysis. Trends in Food Science & Technology, 68, 152–159. 10.1016/j.tifs.2017.08.016

Vogel, R. F., Pavlovic, M., Ehrmann, M. A., Wiezer, A., Liesegang, H., Offschanka, S., Voget, S., Angelov, A., Böcker, G., & Liebl, W. (2011). Genomic analysis reveals Lactobacillus sanfranciscensis as stable element in traditional sourdoughs. Microbial Cell Factories, 10(1), S6. 10.1186/1475-2859-10-S1-S6

Wang, B., Shao, Y., Chen, T., Chen, W., & Chen, F. (2015). Global insights into acetic acid resistance mechanisms and genetic stability of Acetobacter pasteurianus strains by comparative genomics. Scientific Reports, 5(1), 18330. 10.1038/srep18330

Wei, Q., Wang, X., Sun, D.-W., & Pu, H. (2019). Rapid detection and control of psychrotrophic microorganisms in cold storage foods: A review. Trends in Food Science & Technology, 86, 453–464. 10.1016/j.tifs.2019.02.009

