## Supplementary information for "Shipped and shifted: modeling collection-induced bias in microbiome multi-omics using a tractable fermentation system"

### Supplementary figures

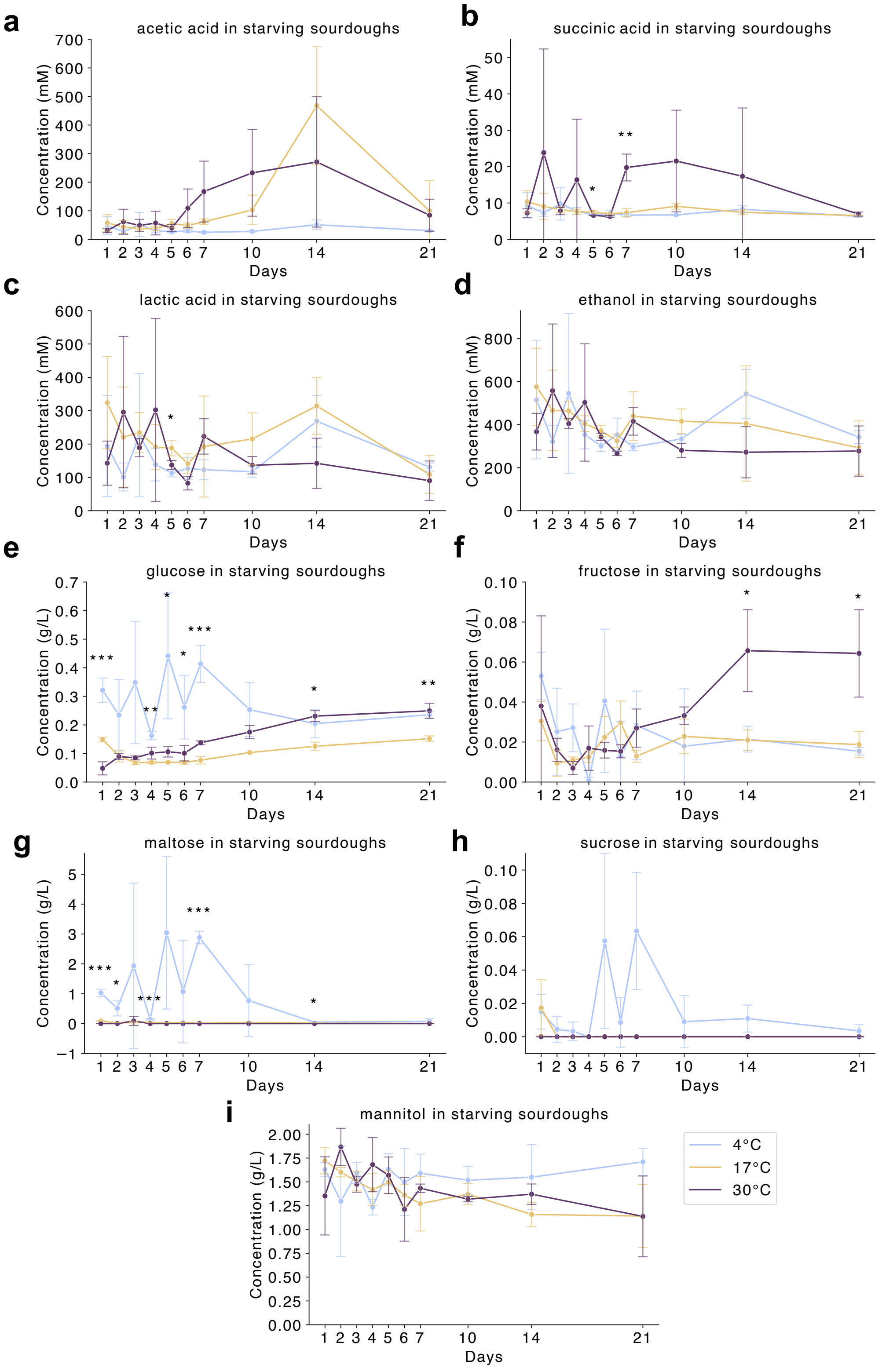


**Supplementary Figure 1. Changes in fermentation-relevant acids and sugars in sourdough samples stored at 4 °C, 17 °C, and 30 °C over 21 days**. Quantified by HPLC, metabolites include acetic acid, succinic acid, lactic acid, ethanol, glucose, fructose, maltose, sucrose and mannitol. Statistical differences between storage temperatures at each time point were determined by one-way ANOVA with FDR correction (*$p < 0.05$, **$p < 0.01$, ***$p < 0.001$).

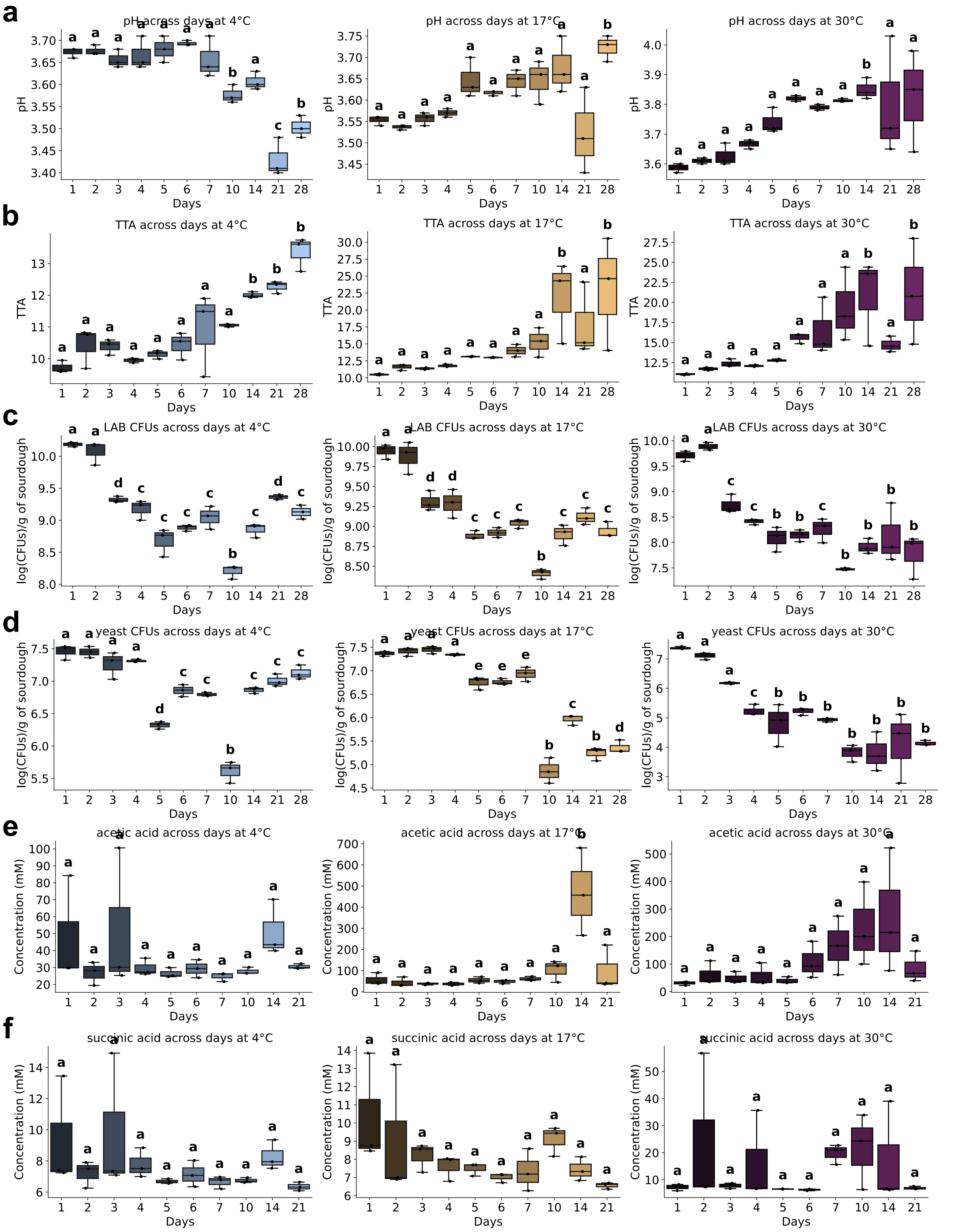


**Supplementary Figure 2. Temporal dynamics of acidity, microbial viability, and organic acid accumulation in sourdough samples stored at 4 °C, 17 °C, and 30 °C**. Measurements over the 28-day storage period are shown for **(a)** pH, **(b)** total titratable acidity (TTA), **(c)** viable lactic acid bacteria (LAB) counts, and **(d)** viable yeast counts; organic acid concentrations were monitored over 21 days for **(e)** acetic acid and **(f)** succinic acid. Statistical comparisons of time points within each temperature group were conducted using post hoc Tukey's HSD with FDR correction. Time points labeled with the same letter are not significantly different (adjusted $p > 0.05$), while those with different letters indicate significant differences (adjusted $p < 0.05$).

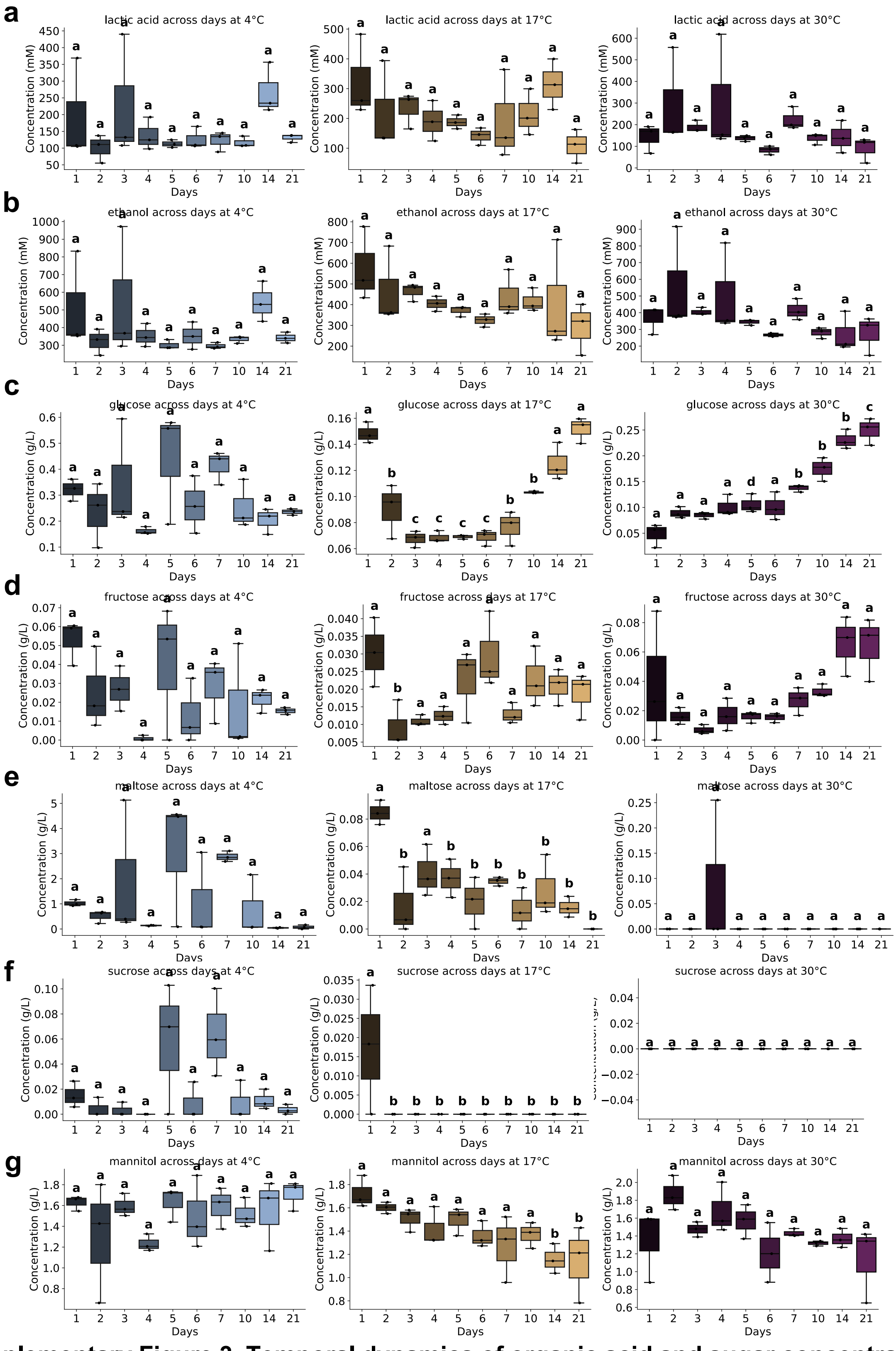


**Supplementary Figure 3. Temporal dynamics of organic acid and sugar concentrations in sourdough samples stored at 4 °C, 17 °C, and 30 °C over a 21-day period.** HPLC-quantified metabolites include: **(a)** lactic acid, **(b)** ethanol, **(c)** glucose, **(d)** fructose, **(e)** maltose, **(f)** sucrose, and **(g)** mannitol. Statistical comparisons of time points within each temperature group were performed using post hoc Tukey's HSD with FDR correction. Time points sharing the same letter are not significantly different (adjusted $p > 0.05$), whereas those with different letters indicate statistically significant differences (adjusted $p < 0.05$).

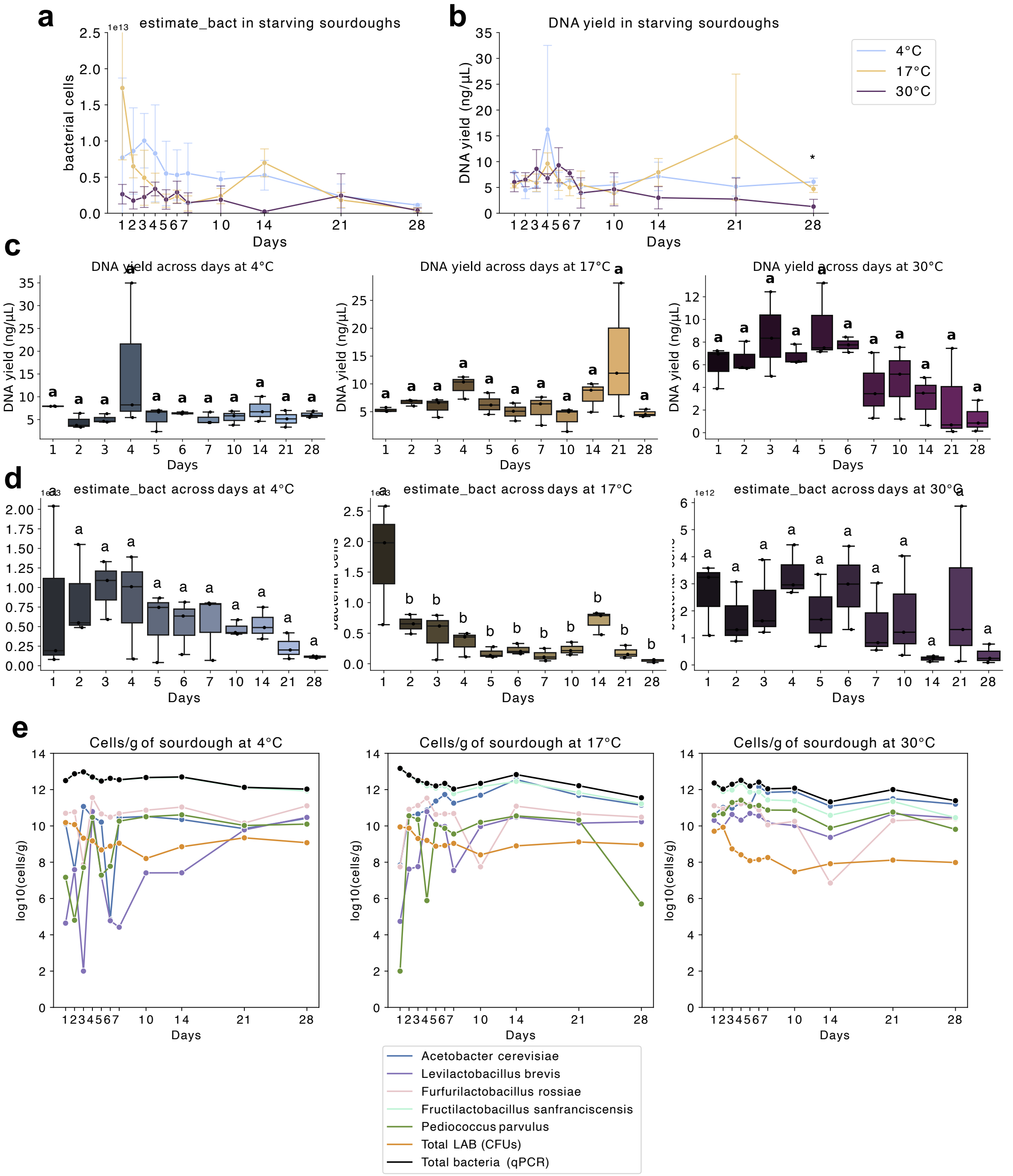


**Supplementary Figure 4. Quantification of total bacterial load and DNA extraction efficiency in sourdough samples stored at 4 °C, 17 °C, and 30 °C over 28 days. a,** Estimated total bacterial cells per gram of sourdough, determined by quantitative PCR (qPCR) targeting the V4 region of the 16S rRNA gene using EMP primers, normalized to the average 16S gene copy number in lactobacilli. **b,** DNA yield (ng/μL) obtained from extractions using the MagMAX Microbiome Ultra kit (Thermo Fisher). Statistical comparisons across temperatures at each time point (panels a, b) were performed using one-way ANOVA with FDR correction (*p < 0.05, **p < 0.01, ***p < 0.001). Temporal comparisons within each temperature group were conducted using post hoc Tukey's HSD with FDR correction for **(c)** DNA yield and **(d)** total bacterial cell estimates. Time points labeled with the same letter are not significantly different; those with different letters indicate significant differences (adjusted $p < 0.05$). **e,** Absolute abundances of dominant bacterial species over time, derived by multiplying 16S rRNA-based relative abundances with qPCR-derived total bacterial estimates. LAB colony-forming units (CFUs) are overlaid for comparative validation of viability versus total bacterial DNA abundance.

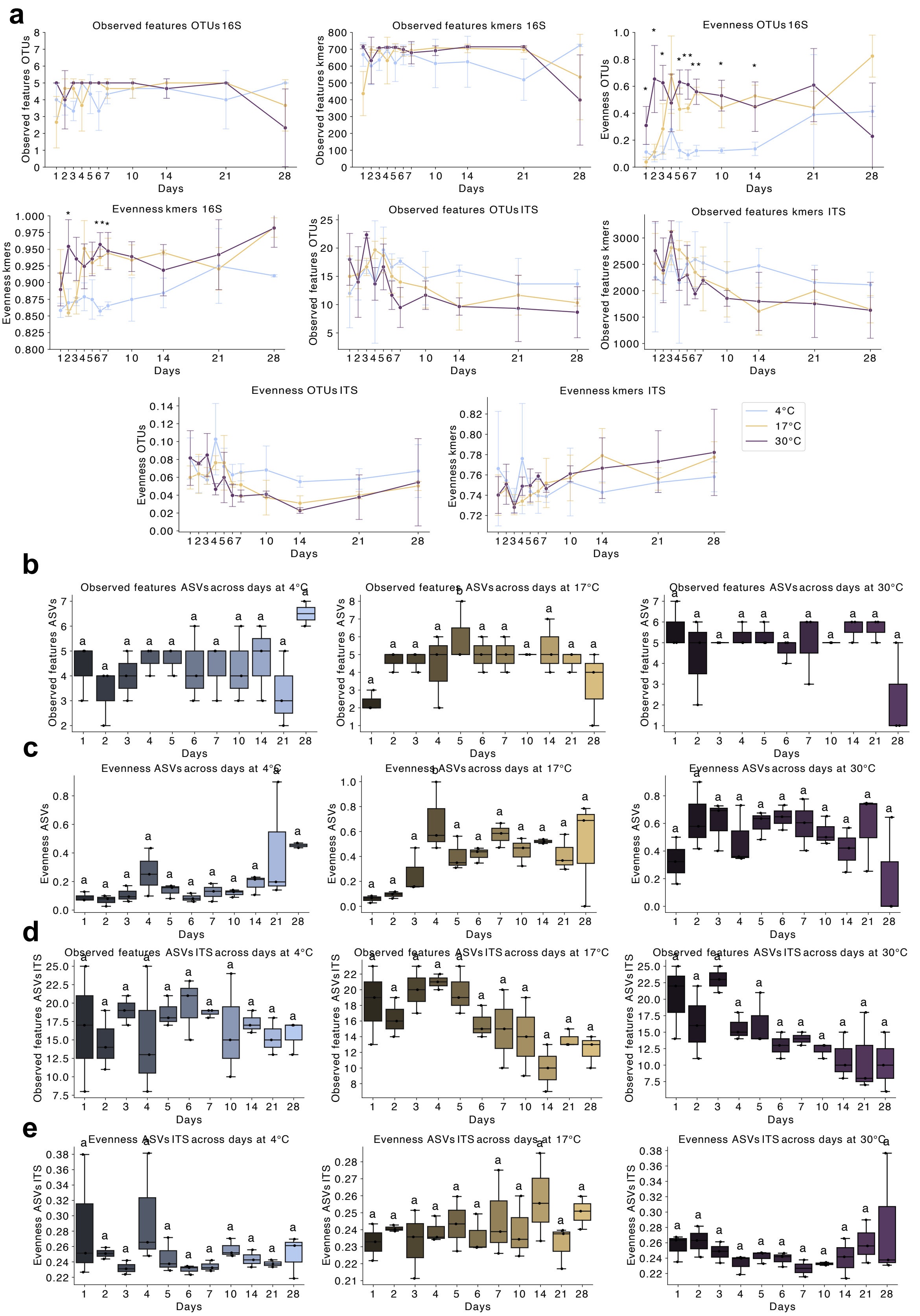


**Supplementary Figure 5. Alpha diversity metrics of bacterial (16S V4) and fungal (ITS) communities in sourdough samples stored at 4 °C, 17 °C, and 30 °C over 28 days. a,** Observed features and community evenness for bacteria and fungi, computed from both OTU clustering and k-mer–based analysis. Statistical comparisons across temperatures at each time point were conducted using one-way ANOVA with FDR correction (*$p < 0.05$, **$p < 0.01$,

***p < 0.001). Temporal comparisons within each temperature group were performed using post hoc Tukey's HSD with FDR correction for: **(b)** bacterial richness (observed ASVs, 16S V4), **(c)** bacterial evenness (ASVs, 16S V4), **(d)** fungal richness (observed ASVs, ITS), and **(e)** fungal evenness (ASVs, ITS). Time points sharing the same letter are not significantly different; those with different letters represent significant changes (adjusted $p < 0.05$).

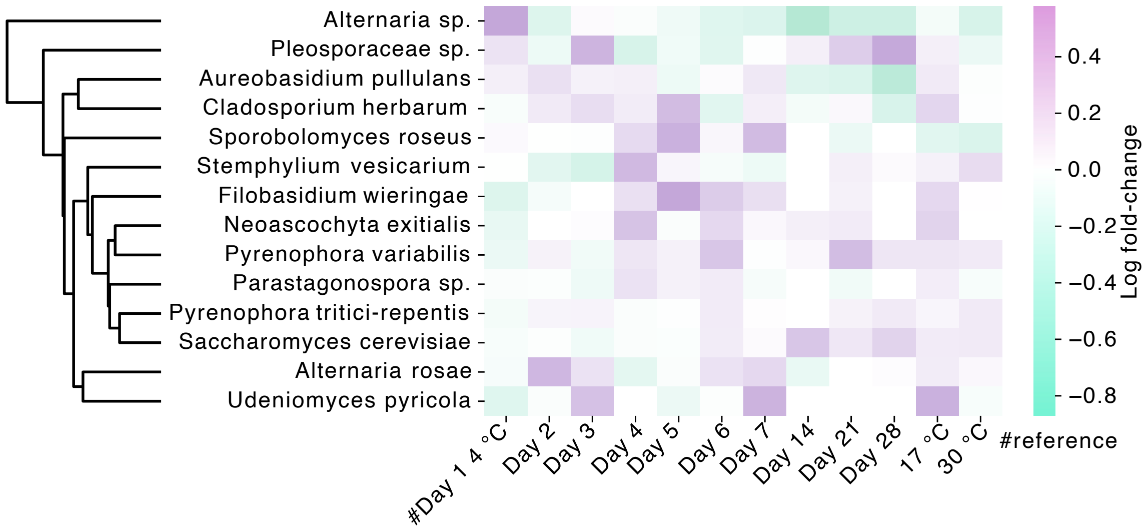


**Supplementary Figure 6. Differential abundance analysis of fungal taxa in sourdough samples stored at 4 °C, 17 °C, and 30 °C over 28 days.** Heatmap of log fold-changes in fungal (ITS) species relative to day 1 at 4 °C, computed using ANCOM-BC2. Taxa were hierarchically clustered on their log fold-changes using average linkage and Euclidean distance. No taxa exhibited statistically significant differential abundance across time or temperature (FDR-adjusted $q < 0.05$).

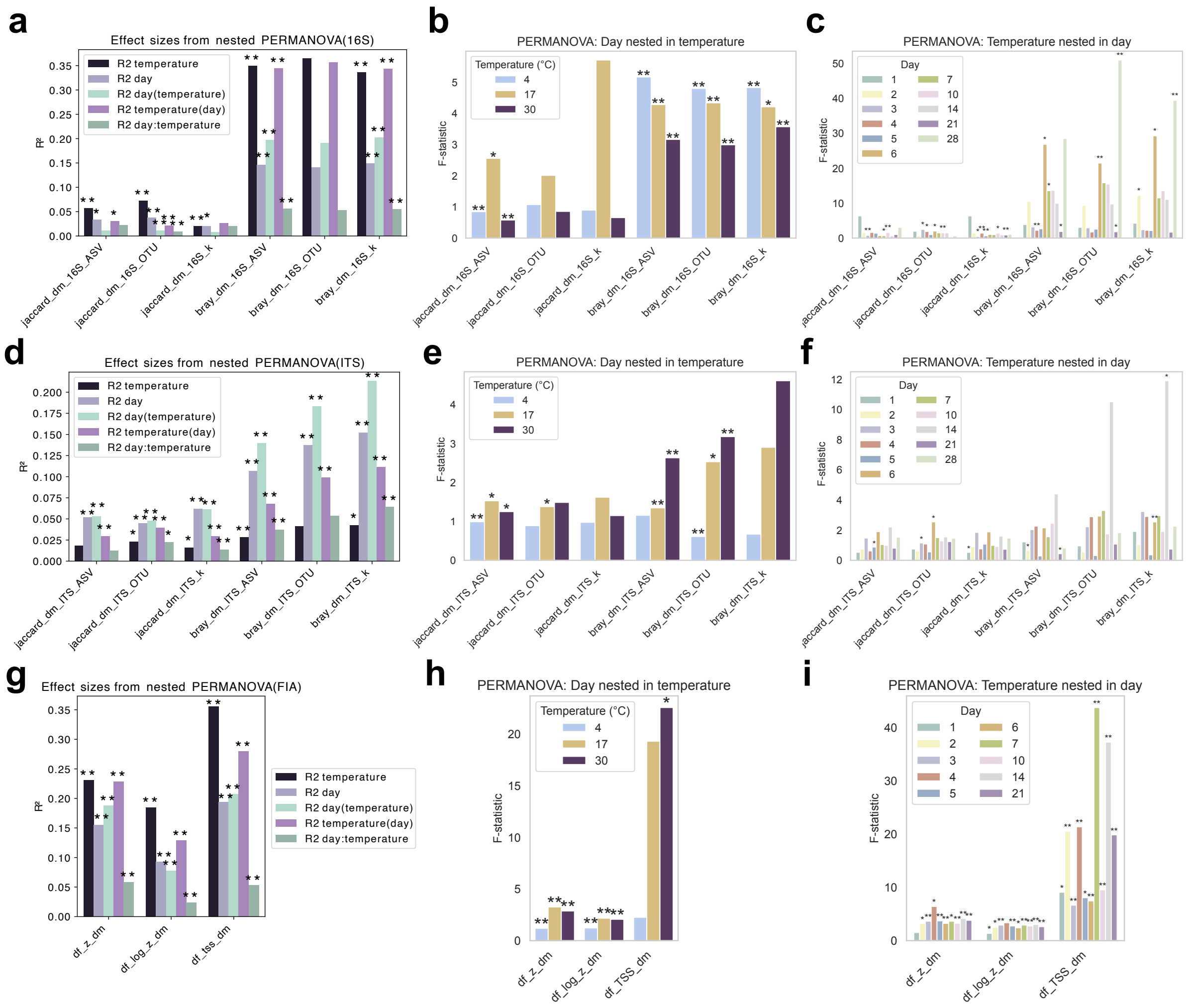


**Supplementary Figure 7. Effect sizes of temperature, time, and their interactions on microbial community structure and metabolomic profiles in sourdough samples**. Variance explained by each factor was assessed using PERMANOVA on distance matrices derived from bacterial and fungal community data (Jaccard and Bray–Curtis, using ASV, OTU, and k-mer feature tables) and from metabolomic profiles (FIA-MS, using z-score, log-z-score, or TSS-transformed data). **a,** Global effects of temperature, day, and their interaction on bacterial beta-diversity. **b,** Fine-grained effects of day within each temperature group on bacterial communities. **c,** Comparisons of temperatures within individual days for bacterial variance. **(d–f)** Equivalent analyses as in a–c, respectively, for fungal beta-diversity. **g,** Effects of temperature, day, and nested variables on global metabolome variance. **h,** Day effects nested within each temperature group, and **(i)** temperature effects nested within individual days for metabolomic variance. Statistical significance was assessed using 999 permutations (*$p < 0.05$, **$p < 0.01$, ***$p < 0.001$).

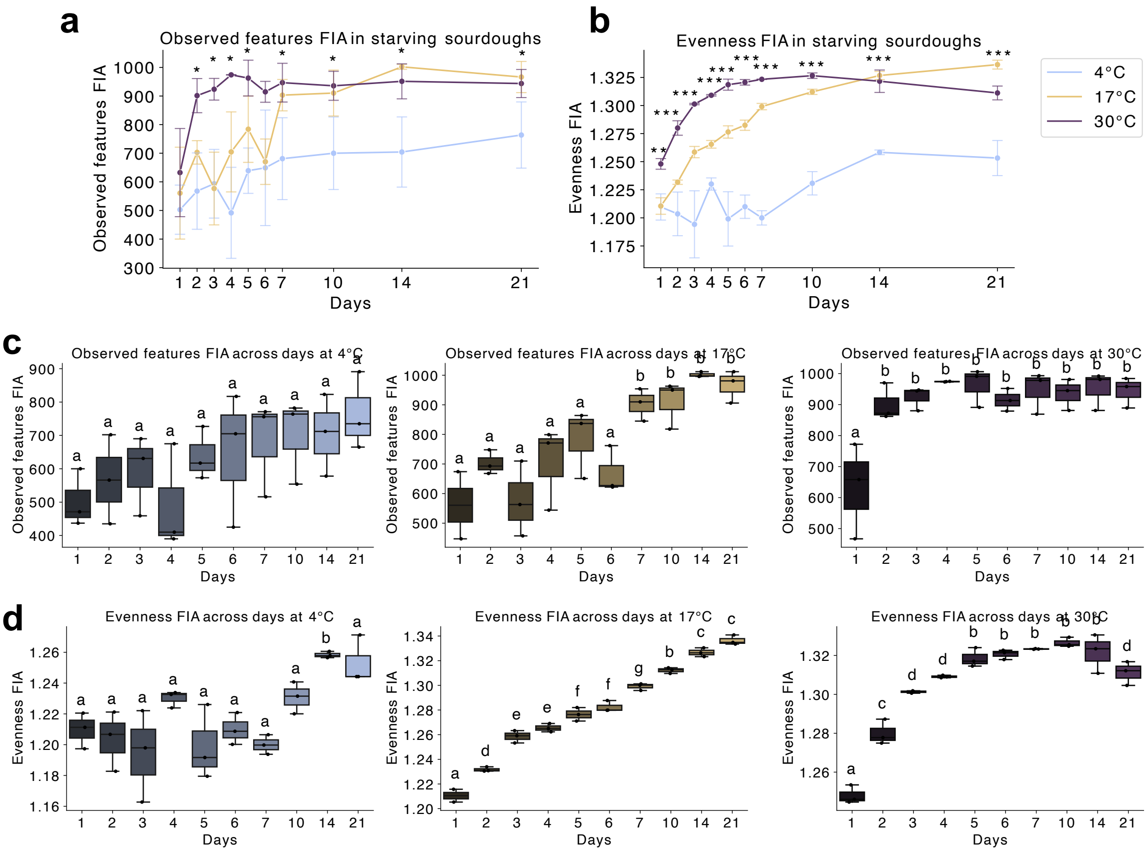


**Supplementary Figure 8. Temporal dynamics of metabolite diversity in sourdough samples stored at 4 °C, 17 °C, and 30 °C over 21 days. a,** Metabolic richness and **(b)** evenness, based on features detected by flow injection analysis–mass spectrometry (FIA-MS). Statistical comparisons across temperature groups at each time point were conducted using one-way ANOVA with FDR correction (*p < 0.05, **p < 0.01, ***p < 0.001). Within-group temporal comparisons were assessed using post hoc Tukey's HSD with FDR correction for: **(c)** observed metabolite features and **(d)** evenness. Time points sharing the same letter are not significantly different; those with different letters indicate statistically significant changes (adjusted $p < 0.05$).

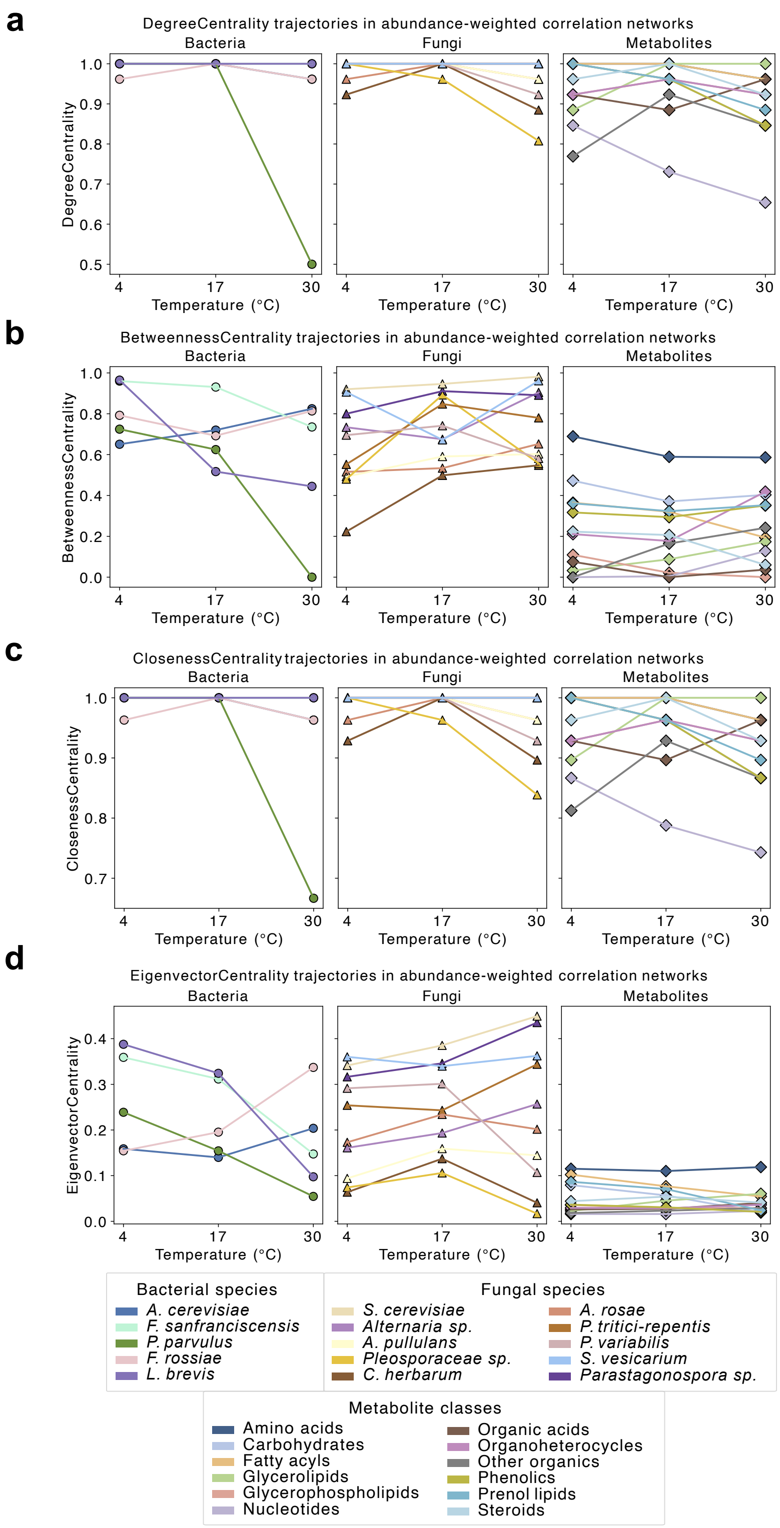


**Supplementary Figure 9. Centrality metrics of abundance-weighted co-correlation networks in sourdough samples stored at 4 °C, 17 °C, and 30 °C.** Network nodes represent bacterial taxa (left panels), fungal taxa (middle panels), and metabolite classes (right panels), with connections derived from correlation-based associations weighted by feature abundance. Centrality was assessed using four metrics: **(a)** degree centrality, **(b)** betweenness centrality, **(c)** closeness centrality, and **(d)** eigenvector centrality. These metrics reflect the relative influence and connectivity of dominant and subdominant microbial and metabolite features across storage conditions.

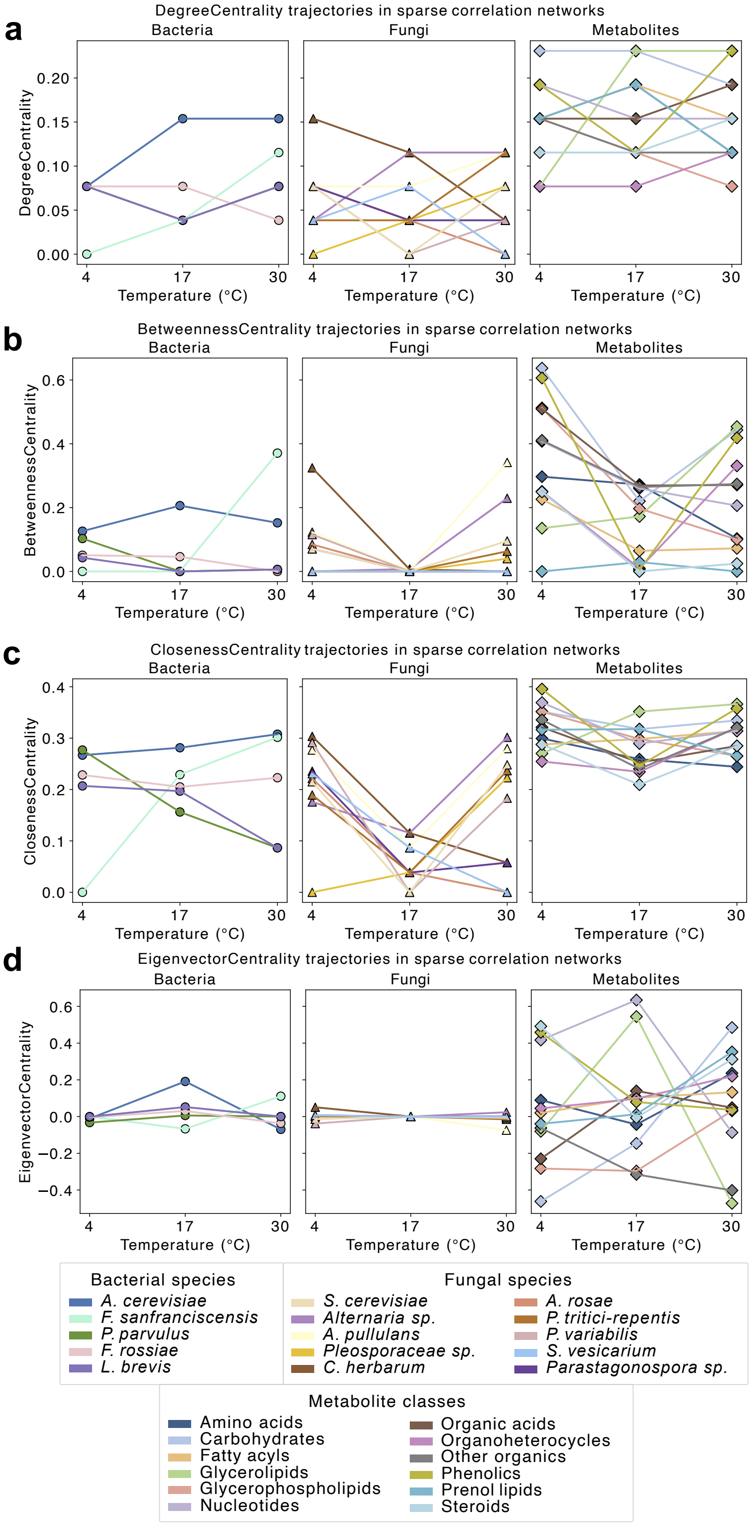


**Supplementary Figure 10. Centrality metrics of sparse correlation networks in sourdough samples stored at 4 °C, 17 °C, and 30 °C**. Network nodes represent bacterial taxa (left panels), fungal taxa (middle panels), and metabolite classes (right panels), with edges derived from sparse inverse covariance estimates using the Graphical Lasso algorithm. Centrality was evaluated using: **(a)** degree centrality, **(b)** betweenness centrality, **(c)** closeness centrality, and **(d)** eigenvector centrality. These metrics characterize the structural connectivity and influence of microbial and metabolite features, independent of direct abundance co-correlations, across different storage conditions.

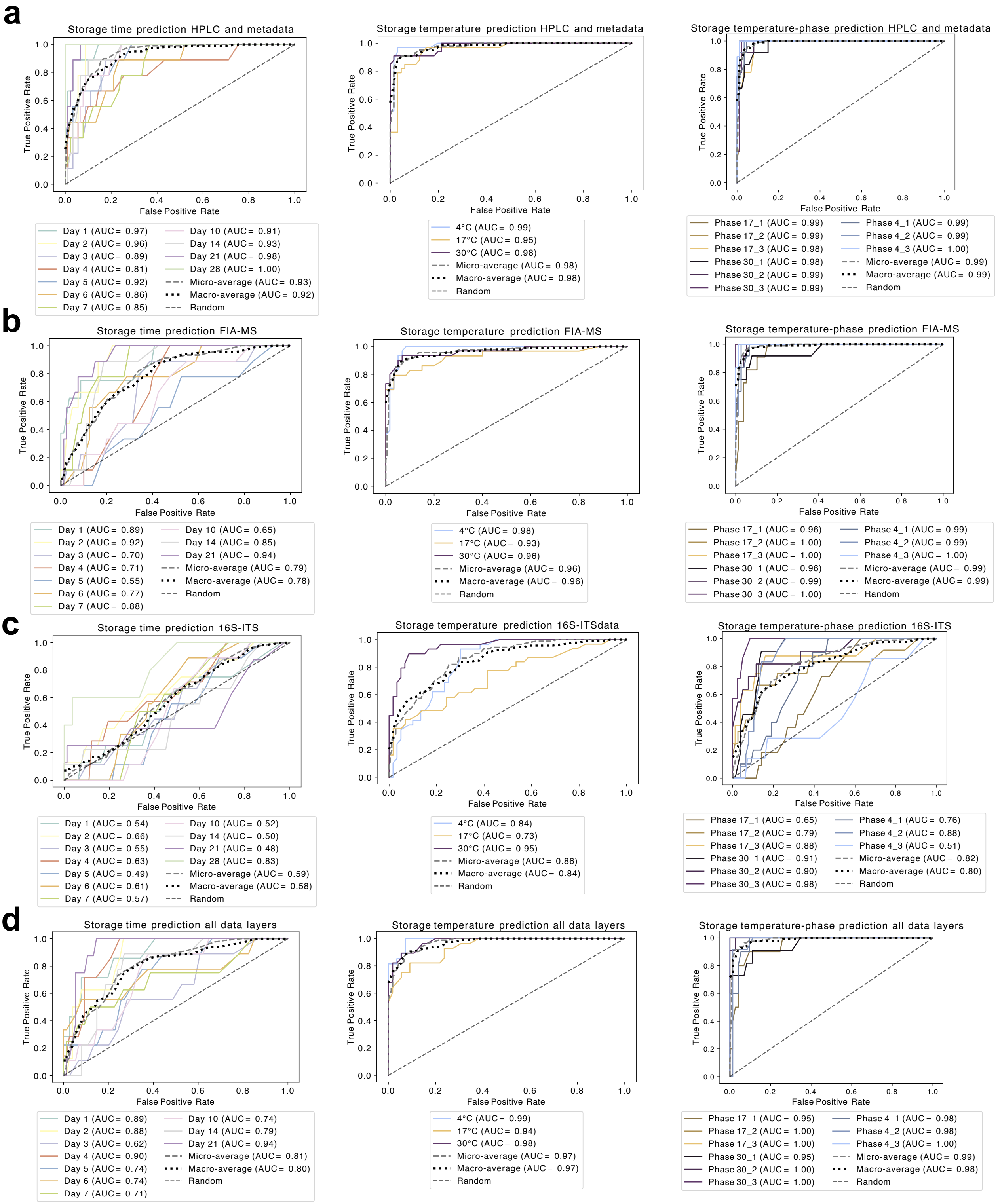


**Supplementary Figure 11. Predictive accuracy of storage parameters based on multi-omics and physicochemical data layers.** Classification performance of random forest models (100 estimators, 5-fold cross-validation) is reported as area under the curve (AUC) for predicting storage day (left), storage temperature (middle), and combined storage phase (right; Phase 1: days 1–4, Phase 2: days 5–10, Phase 3: days 14–28). Models were trained using: **(a)** HPLC-derived sugar and acid profiles combined with colony-forming units (CFU), pH, and total titratable acidity (TTA); **(b)** metabolite features from FIA-MS; **(c)** CLR-transformed relative abundances from 16S V4 rRNA and ITS amplicon sequencing; and **(d)** a combined model integrating all features from (a), (b), and (c).

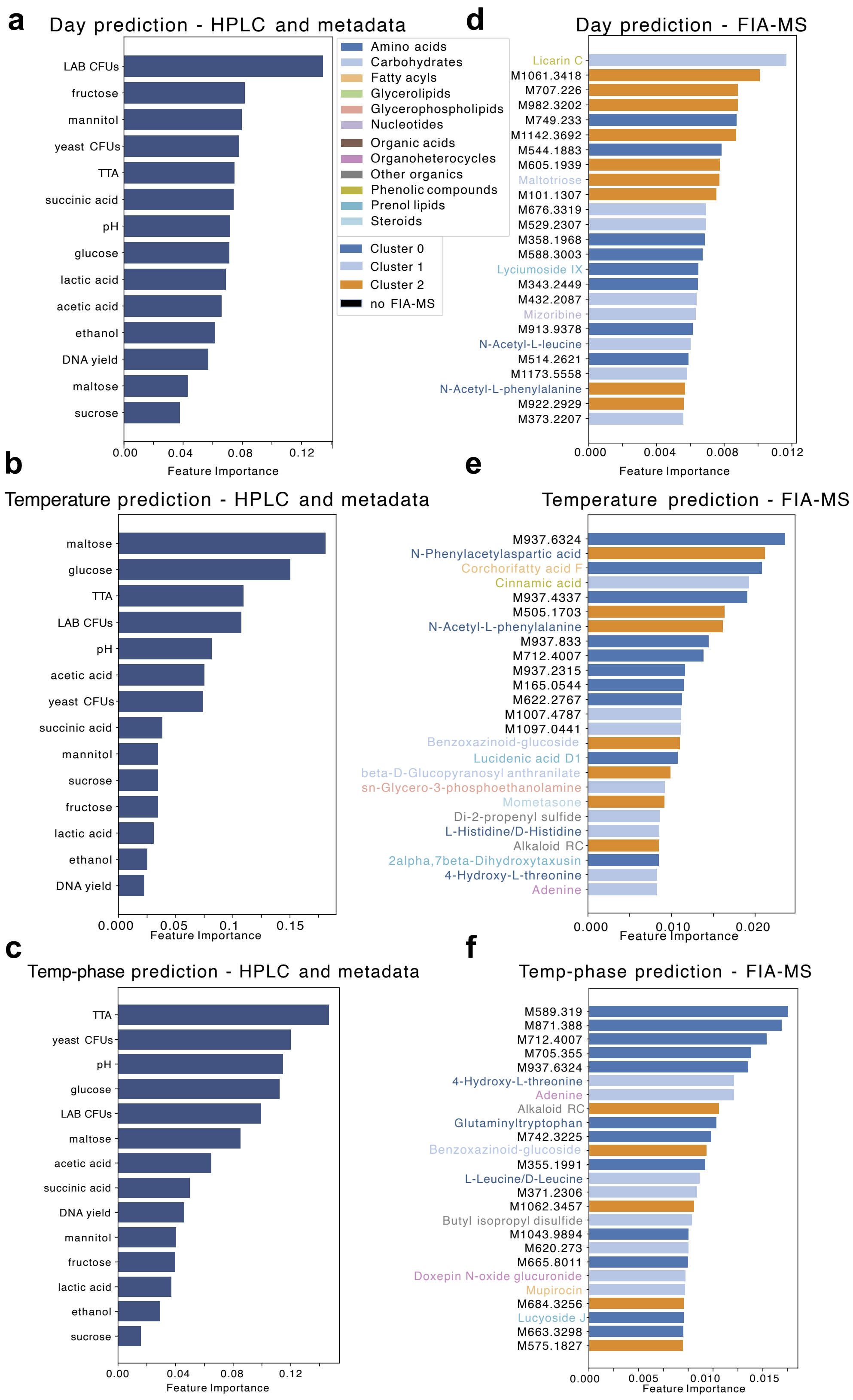


**Supplementary Figure 12. Top predictive features for classifying storage parameters based on HPLC-metadata and FIA-MS data.** Feature importance scores from random forest models are shown for predicting (top) storage day, (middle) storage temperature, and (bottom) combined storage phase (Phase 1: days 1–4, Phase 2: days 5–10, Phase 3: days 14–28), using: **(a–c)** HPLC-derived sugar and acid concentrations, CFU counts, pH, and total titratable acidity (TTA); and **(d–f)** metabolite features detected via FIA-MS. FIA-MS features were annotated where m/z values matched entries in the KEGG or HMDB databases within a mass tolerance of 0.002 Da, bars are colored according to the major dynamic metabolite clusters over time, and labels based on metabolite class. Models were trained with 100 estimators and 5-fold cross-validation. Top-ranked features reflect the most informative signals associated with temporal and temperature-driven changes in sourdough fermentation profiles.

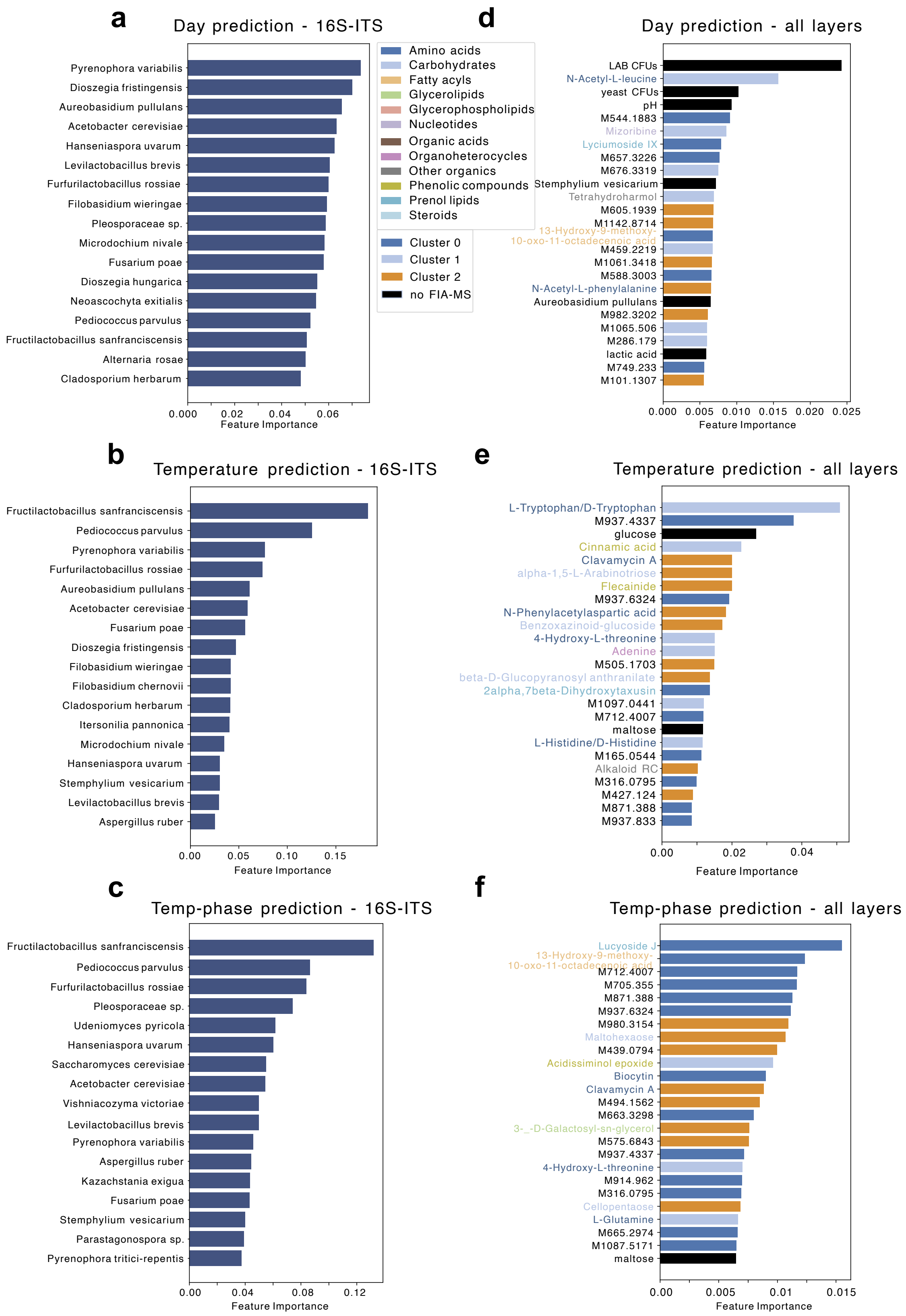

**Supplementary Figure 13. Top predictive features for classifying storage parameters based on microbiome and fully integrated multi-omics data.** Feature importance scores from random forest models are shown for predicting (top) storage day, (middle) storage temperature, and (bottom) combined storage phase per temperature (Phase 1: days 1–4, Phase 2: days 5–10, Phase 3: days 14–28), using: **(a–c)** microbiome features from combined bacterial (16S V4) and fungal (ITS) amplicon profiles; and **(d–f)** an integrated dataset comprising microbiome, metabolome (FIA-MS and HPLC), CFU counts, pH, and total titratable acidity (TTA). Bars are colored according to the major dynamic FIA-MS metabolite clusters over time, and labels based on FIA-MS metabolite class. All models were trained with 100 estimators and 5-fold cross-validation. Top-ranked features represent the most informative variables contributing to the discrimination of temporal and temperature-dependent sourdough changes.

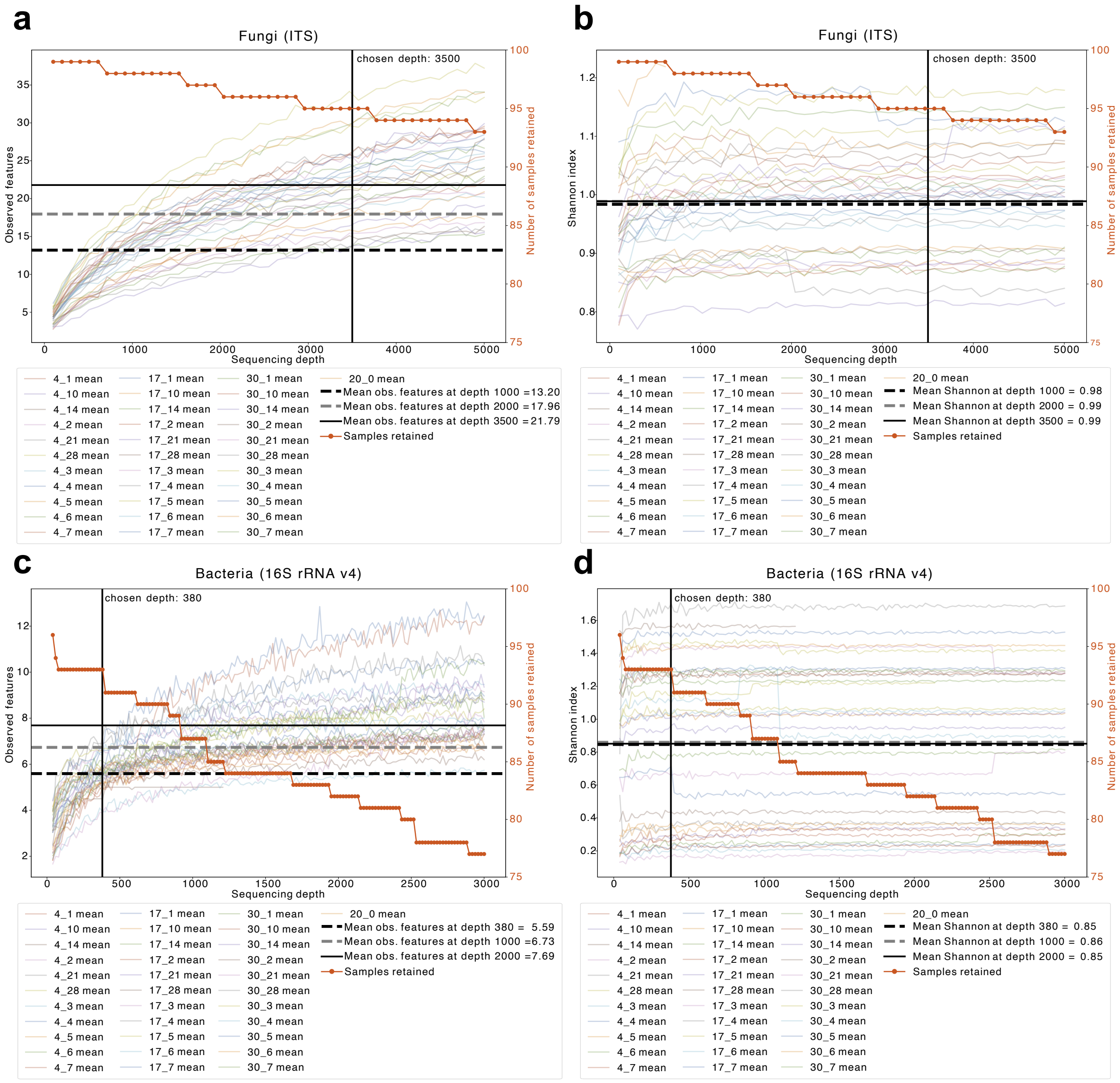


**Supplementary Figure 14. Alpha-rarefaction curves and sample retention profiles for fungal and bacterial communities. (a, b)** Fungal ASVs: rarefaction curves overlaid with sample retention curves for (a) observed features and (b) Shannon diversity; the selected sequencing depth of 3500 is indicated. **(c, d)** Bacterial ASVs: corresponding overlays for (c) observed features and (d) Shannon diversity, highlighting a chosen depth of 380 as the optimal trade-off between diversity representation and sample retention.

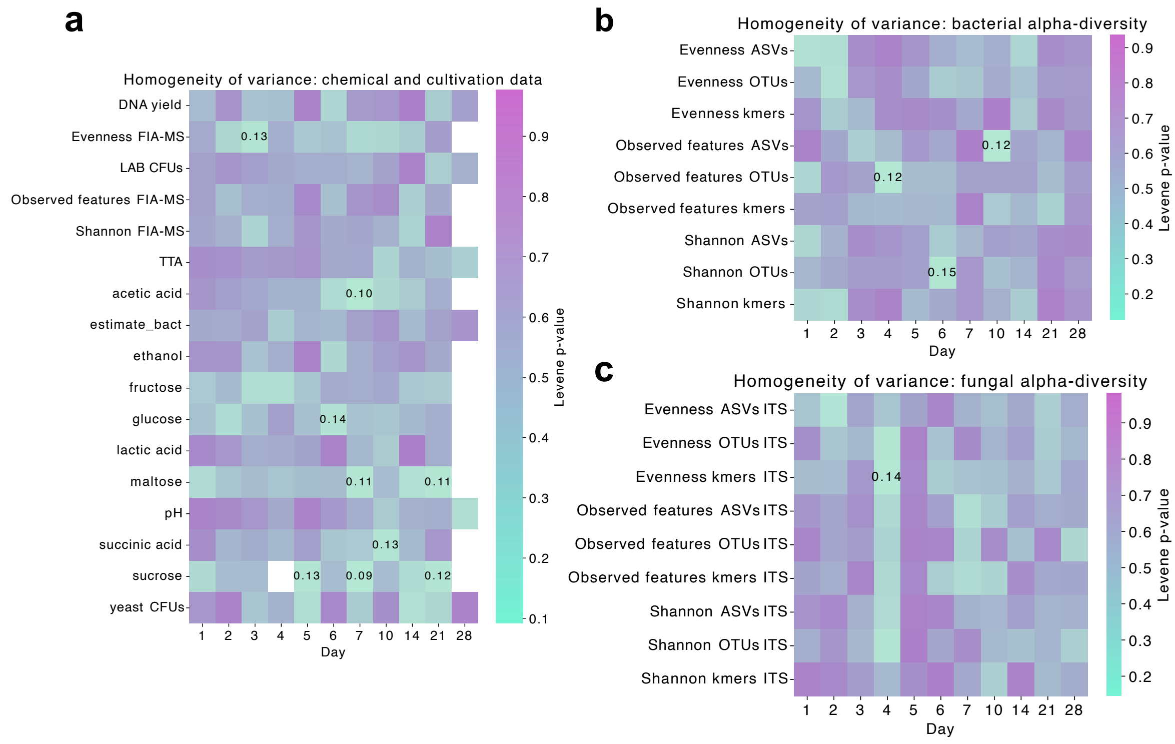


**Supplementary Figure 15. Homogeneity of variance evaluation across temperatures.** Heatmaps showing Levene's test p-values for comparing variance across storage temperatures (4 °C, 17 °C, and 30 °C) per day and metric. **a,** Chemical and cultivation-based data (HPLC, FIA-MS alpha-diversity, CFUs, DNA yield, and qPCR-based bacterial load estimate); **(b)**, bacterial alpha-diversity metrics; **(c)**, fungal alpha-diversity metrics across ASV, OTU, and k-mer resolutions. This analysis was conducted to evaluate the assumption of equal variances prior to performing ANOVA at each timepoint. P-values < 0.15 are annotated, yet none fall below the conventional threshold of 0.05, indicating no statistically significant heterogeneity of variance between temperature groups. These results support the suitability of ANOVA for downstream comparisons, even under small sample sizes. Chemical data (HPLC and FIA-MS) was not acquired for day 28 samples, therefore no p-values are displayed in (a) for those metrics at day 28.

### Supplementary tables

**Supplementary Table 1. Composition and 16S rRNA gene copy number normalization of the ZymoBIOMICS Microbial Community DNA standard.** This table lists microbial species, relative DNA abundance, genome size, ploidy, 16S/18S rRNA gene copy number, genome mass, and the calculated number of genomes and 16S genes per µL assuming 100 ng total DNA input. All values for genome size, ploidy, copy number, and genome mass were obtained from the ZymoBIOMICS Microbial Community DNA Standard (Cat. No. D6305) reference data. The total bacterial 16S rRNA gene abundance per µL with 100 ng total DNA was calculated as $1.58 \times 10^{11}$ copies, corresponding to $2.65 \times 10^{10}$ bacterial genomes. A mean normalization factor of 5.95 16S rRNA gene copies per genome can be derived by dividing total 16S abundance by total genome number ($1.58 \times 10^{11} / 2.65 \times 10^{10}$), a fixed factor of 5 was applied in this study, reflecting typical 16S copy numbers in dominant taxa such as the *Lactobacillus* representative in the standard, *Lactobacillus fermentum*. This correction factor was used to normalize 16S qPCR-based bacterial load estimates in sourdough samples, together with the assumption of $1.58 \times 10^{9}$ gene copies per ng DNA.

| Species | % DNA | ng DNA per 1 uL | genome size (Mb) | ploidy | 16S/18S copy number | genome mass (in ng) | genomes per uL (100 ng) | 16S genes per 1 uL with 100 ng DNA |
|---|---|---|---|---|---|---|---|---|
| *Listeria monocytogenes* | 0.12 | 12 | 2992 | 1 | 6 | 3.23E-09 | 3715754833 | 22294529000 |
| *Pseudonomas aeruginosa* | 0.12 | 12 | 6792 | 1 | 4 | 7.33E-09 | 1636857842 | 6547431367 |
| *Bacillus subtilis* | 0.12 | 12 | 4045 | 1 | 10 | 4.37E-09 | 2748464391 | 27484643910 |
| *Escherichia coli* | 0.12 | 12 | 4875 | 1 | 7 | 5.26E-09 | 2280520710 | 15963644970 |
| *Salmonella enterica* | 0.12 | 12 | 4760 | 1 | 7 | 5.14E-09 | 2335617324 | 16349321267 |
| *Lactobacillus fermentum* | 0.12 | 12 | 1905 | 1 | 5 | 2.06E-09 | 5835978195 | 29179890975 |
| *Enterococcus faecalis* | 0.12 | 12 | 2845 | 1 | 4 | 3.07E-09 | 3907746384 | 15630985535 |
| *Staphylococcus aureus* | 0.12 | 12 | 2730 | 1 | 6 | 2.95E-09 | 4072358411 | 24434150465 |
| *Saccharomyces cerevisiae* | 0.02 | 2 | 12100 | 2 | 109 | 1.31E-08 | 153134138.6 | 16691621106 |
| *Cryptococcus neoformans* | 0.02 | 2 | 18900 | 2 | 60 | 2.04E-08 | 98038258.04 | 5882295482 |

#### Supplementary Table 2. Gradient used for Carbohydrate separation by HPLC.

| *Time (min.)* | *Flow rate (mL/min.)* | *Mobile phase 1 in %* | *Mobile phase 2 in %* | *Mobile phase 3 in %* |
|---|---|---|---|---|
| *0* | 0.25 | 35 | 0.2 | 64.8 |
| *5* | 0.25 | 35 | 0.2 | 64.8 |
| *20* | 0.25 | 35 | 3 | 62 |
| *30* | 0.25 | 60 | 30 | 10 |
| *35* | 0.25 | 35 | 0.2 | 64.8 |
| *45* | 0.25 | 35 | 0.2 | 64.8 |

**Supplementary Table 3. Differentially abundant bacterial taxa in the reference group.** ANCOM-BC2 identified taxa enriched in sourdough samples stored at 4 °C on day 1. For log-ratio analyses over time (see Fig. 2e), only taxa with absolute log-fold changes (|logFC|) > 0.75 were retained. Based on the enrichments observed here, *Acetobacter cerevisiae* was selected as the denominator taxon, while *Fructilactobacillus sanfranciscensis* and *Furfurilactobacillus rossiae* were used as numerator taxa for all subsequent log-ratio calculations across time points.

| *taxon* | *logFC* | *qval* | *pval* |
|---|---|---|---|
| *Acetobacter cerevisiae* | -1.6052 | 0.97919 | 0.493079 |
| *Fructilactobacillus sanfranciscensis* | 2.588816 | 0.97919 | 0.231429 |
| *Pediococcus parvulus* | -0.06316 | 0.97919 | 0.97919 |
| *Furfurilactobacillus rossiae* | 0.94978 | 0.97919 | 0.666257 |
| *Levilactobacillus brevis* | 0.622505 | 0.97919 | 0.805273 |

**Supplementary Table 4. Differentially abundant fungal taxa in the reference group.** ANCOM-BC2 identified taxa enriched in sourdough samples stored at 4 °C on day 1. For log-ratio analyses over time (see Fig. 2f), only taxa with absolute log-fold changes (|logFC|) > 0.1 were retained. Based on the enrichments observed here, *Filobasidium wieringae, Neoascochyta exitialis, Pyrenophora tritici-repentis, Pyrenophora variabilis* and *Udeniomyces pyricola* were selected as the denominator taxa, while *Alternaria sp.* and *Pleosporaceae sp.* were used as numerator taxa for all subsequent log-ratio calculations across time points.

| *taxon* | *logFC* | *qval* | *pval* |
|---|---|---|---|
| *Alternaria rosae* | -0.085088 | 0.997784 | 0.98185 |
| *Alternaria sp.* | 0.578727 | 0.997784 | 0.852397 |
| *Aureobasidium pullulans* | 0.093354 | 0.997784 | 0.977036 |
| *Cladosporium herbarum* | -0.068014 | 0.997784 | 0.984249 |
| *Filobasidium wieringae* | -0.349554 | 0.997784 | 0.943255 |
| *Neoascochyta exitialis* | -0.230582 | 0.997784 | 0.961077 |
| *Parastagonospora sp.* | -0.053159 | 0.997784 | 0.990194 |
| *Pleosporaceae sp.* | 0.179644 | 0.997784 | 0.9592 |
| *Pyrenophora tritici-repentis* | -0.107098 | 0.997784 | 0.977447 |
| *Pyrenophora variabilis* | -0.19592 | 0.997784 | 0.959906 |
| *Saccharomyces cerevisiae* | -0.075105 | 0.997784 | 0.98019 |
| *Sporobolomyces roseus* | 0.036263 | 0.997784 | 0.994492 |
| *Stemphylium vesicarium* | -0.011866 | 0.997784 | 0.997784 |
| *Udeniomyces pyricola* | -0.325637 | 0.997784 | 0.956649 |

**Supplementary Table 5. Concordance between bacterial community composition and metabolome profiles.** Procrustes analysis comparing 16S v4 rRNA gene-based Bray–Curtis distances with various transformed and dimensionally reduced FIA-MS metabolomics datasets.

| *comparison* | *Observed Disparity* | *95% CI* | *pval* |
|---|---|---|---|
| *FIA_z_PCoA vs 16S_PCoA* | 0.591 | [0.953632, 0.998986] | 0.001 |
| *FIA_tss_PCoA vs 16S_PCoA* | 0.437 | [0.946777, 0.999179] | 0.001 |
| *FIA_tss_PCA vs 16S_PCoA* | 0.499 | [0.944602, 0.998966] | 0.001 |
| *FIA_z_PCA vs 16S_PCoA* | 0.591 | [0.947394, 0.998983] | 0.001 |

**Supplementary Table 6. Temporal concordance between bacterial communities and metabolome profiles.** Procrustes analysis comparing Bray–Curtis distances from 16S V4 rRNA gene data and TSS-normalized FIA-MS metabolomics profiles, both projected into PCoA space across time points.

| *day* | *disparity* | *pval* |
|---|---|---|
| *1* | 0.5755 | 0.0350 |
| *2* | 0.3473 | 0.0080 |
| *3* | 0.5785 | 0.0180 |
| *4* | 0.7841 | 0.3107 |
| *5* | 0.6087 | 0.0380 |
| *6* | 0.2776 | 0.0020 |
| *7* | 0.2416 | 0.0030 |
| *10* | 0.2974 | 0.0050 |
| *14* | 0.2552 | 0.0040 |
| *21* | 0.5397 | 0.0539 |

**Supplementary Table 7. Temporal concordance by temperature between bacterial communities and metabolome profiles.** Procrustes analysis comparing Bray–Curtis distances from 16S V4 rRNA gene data and TSS-normalized FIA-MS metabolomics profiles, both projected into PCoA space across time points per temperature.

| *day* | *temperature* | *disparity* | *pval* |
|---|---|---|---|
| *1* | 4 | 0.938 | 1.000 |
| *2* | 4 | 0.520 | 0.677 |
| *3* | 4 | 0.797 | 0.689 |
| *4* | 4 | 0.090 | 0.001 |
| *5* | 4 | 0.014 | 0.191 |
| *6* | 4 | 0.534 | 0.341 |
| *7* | 4 | 0.616 | 0.495 |
| *10* | 4 | 0.117 | 0.169 |
| *14* | 4 | 0.042 | 0.316 |
| *21* | 4 | 0.000 | 0.001 |
| *1* | 17 | 0.000 | 0.001 |
| *2* | 17 | 0.472 | 0.479 |
| *3* | 17 | 0.046 | 0.001 |
| *4* | 17 | 0.000 | 1.000 |
| *5* | 17 | 0.248 | 0.353 |
| *6* | 17 | 0.310 | 0.371 |
| *7* | 17 | 0.247 | 0.518 |
| *10* | 17 | 0.950 | 1.000 |
| *14* | 17 | 0.285 | 0.515 |
| *21* | 17 | 0.023 | 0.155 |
| *1* | 30 | 0.848 | 0.687 |
| *2* | 30 | 0.000 | 1.000 |
| *3* | 30 | 0.266 | 0.154 |
| *4* | 30 | 0.749 | 0.683 |
| *5* | 30 | 0.389 | 0.543 |
| *6* | 30 | 0.247 | 0.656 |
| *7* | 30 | 0.324 | 0.510 |
| *10* | 30 | 0.591 | 0.836 |
| *14* | 30 | 0.808 | 0.685 |
| *21* | 30 | 0.123 | 0.170 |
